# FMR1 Perturbations Disrupt Sensory Corticostriatal Circuits and Mitochondrial Energetics in Fragile X Syndrome

**DOI:** 10.64898/2026.10.07.756495

**Authors:** Yu Yan, Dongyun Li, Jianyu Yuan, Danyong Feng, Jiarui Chang, Fengyu Zhang, Man Xiong, Yan Jiang, Biao Yan, Qiong Xu, Rihui Li, Yi He, Peipei Liu, Mingshuo Shao, Minjie Shen

**Author notes:** Responsible party and corresponding author: Minjie Shen, Institutes of Brain Science, Fudan University, 131 Dong’an Road, Shanghai 200032, China. Additional corresponding authors: Mingshuo Shao, Peipei Liu, Rihui Li, and Yi He. These authors contributed equally (Yu Yan, Dongyun Li, Jianyu Yuan, Danyong Feng).

## Abstract

Fragile X syndrome (FXS) features sensory, social, and cognitive dysfunction, but how *fragile X messenger ribonucleoprotein 1 (FMR1) deficiency* disrupts sensory corticostriatal circuitry and whether distinct *FMR1* perturbations converge on shared cellular mechanisms remain incompletely defined. We integrated diffusion magnetic resonance imaging (MRI) in individuals with FXS and *Fmr1-knockout (KO) mice* with projection-defined calcium recording, bidirectional optogenetic manipulation, primary somatosensory cortex (S1)-targeted expression of three patient-associated *FMR1* missense variants, regional transcriptomics, and optical manipulation of mitochondrial energetic state. Individuals with FXS exhibited disrupted sensory corticostriatal structural connectivity, with mouse analyses further localizing abnormalities to S1–striatal pathways. *Fmr1* deficiency impaired recruitment of the S1–dorsomedial striatum (DMS) pathway during social interaction, and projection-specific manipulation demonstrated that S1–DMS dysfunction contributes to social and cognitive deficits. Three patient-associated *FMR1* variants, I234T, E329G, and R586Q, converged on social behavioral deficits, which were improved by S1–DMS activation. Transcriptomic analyses revealed convergent oxidative phosphorylation (OXPHOS)-related mitochondrial remodeling in S1 and DMS across *Fmr1* deficiency and variant models. Systemic treatment with the mitochondria-targeted antioxidant MitoQ attenuated novel-location-recognition, social-interaction, and social-novelty deficits in *Fmr1*-KO mice. Moreover, mt-EcGAPR-based Mitochondrial OptoEnergetic Activation (Mito-OptoEA) enhanced mitochondrial energetic state in S1 and improved novel-location recognition and sociability in both *Fmr1*-KO and I234T models. Together, these findings define a sensory corticostriatal circuit–metabolic framework for FXS in which S1–DMS circuit dysfunction and convergent mitochondrial energetic abnormalities contribute to *FMR1*-related social and cognitive phenotypes.

## Introduction

Fragile X syndrome (FXS) is the most common inherited monogenic cause of autism spectrum disorder and is characterized by prominent sensory, social and cognitive dysfunction. In most affected individuals, expansion of the cytosine–guanine–guanine (CGG) repeat in fragile X messenger ribonucleoprotein 1 (*FMR1*) induces methylation-dependent transcriptional silencing and loss of fragile X messenger ribonucleoprotein (FMRP). [1, 2] FMRP regulates the translation of synaptic and autism-associated transcripts [3, 4] and participates more broadly in neuronal RNA metabolism. [5] *Fmr1*-deficient mice show altered synaptic plasticity, [6] while genetic intervention can ameliorate multiple synaptic and behavioral abnormalities. [7] Abnormal neocortical development and barrel-cortex plasticity, [8, 9] reduced tonic inhibition, [10] impaired sensorimotor gating, [11] altered auditory cortical ensembles [12] and a visual-cortical electrophysiological signature shared by individuals with FXS and *Fmr1*-deficient mice [13] collectively implicate distributed sensory-network dysfunction. How these sensory cortical abnormalities propagate through downstream circuits to shape social and cognitive behavior remains unresolved.

Corticostriatal pathways provide a plausible route through which altered sensory cortical activity may influence behavior. Their topographic organization allows cortical representations to influence striatal computation, action selection and disease-relevant behavior, [14, 15] while functionally distinct striatal territories integrate diverse excitatory inputs. [16] Projection-level activity can encode social interaction, [17] and genetic disruption of striatal circuitry or optogenetic modulation of a corticostriatal projection can alter autism-related and compulsive phenotypes. [18, 19] Within this architecture, the projection from primary somatosensory cortex (S1) to dorsomedial striatum (DMS) represents a tractable candidate through which abnormal somatosensory signals could influence social behavior and spatial recognition. The contribution of the S1–DMS pathway to these behavioral abnormalities in FXS, however, remains undefined.

Human diffusion magnetic resonance imaging (MRI) provides a clinically accessible approach to characterize structural abnormalities associated with FXS. Previous studies have identified white-matter microstructural abnormalities, including frontostriatal changes that emerge early in development, [20, 21] altered structural-network segregation [22] and atypical white-matter development in infancy, [23] with tract-level effects varying in direction and distribution. [24] Recent circuit studies demonstrate that cell subtype and inhibitory balance sharpen the mechanistic interpretation of social behavior, [25, 26] while adeno-associated virus (AAV)-enabled multimodal mapping illustrates how structural and functional network measures can be experimentally linked. [27] A cross-species strategy therefore provides a means to identify corticostriatal structural abnormalities in individuals with FXS, localize corresponding pathways in a genetic mouse model, and test their functional contribution to behavior.

Allelic heterogeneity adds a second level of mechanistic complexity. Patient-associated *FMR1* missense substitutions can produce domain-dependent changes in FMRP function, synaptic plasticity and socio-cognitive behavior, [28, 29] and C-terminal mutations further disrupt FMRP localization and function. [30] A clinically described missense mutation shows that an FXS phenotype can arise from a discrete amino-acid substitution rather than the canonical repeat-expansion mechanism, [31] while *FMR1*-mutant marmosets extend allele-dependent phenotyping to a primate model. [32] Despite this genetic diversity, distinct *FMR1* perturbations may converge on shared cellular vulnerabilities. Mitochondrial dysfunction is increasingly implicated in FXS: patient-derived cells exhibit abnormal energetic metabolism, including adenosine triphosphate (ATP)-synthase proton leak [33] and impaired mitochondrial biogenesis, [34] while FMRP regulates mitochondrial dynamics by promoting both mitochondrial fusion [35] and fission [36] through distinct molecular mechanisms. These findings raise the possibility that mitochondrial energetic dysregulation represents a convergent downstream consequence of disrupted FMRP function and a potentially tractable point of intervention. Recent advances in engineered light-driven mitochondrial proton pumps enable direct manipulation of mitochondrial energetic state, [37, 38] providing an experimental approach to test whether mitochondrial energetic remodeling represents a modifiable mechanism across distinct *FMR1* perturbations.

Here, we combined human and mouse diffusion MRI to define sensory corticostriatal structural abnormalities across species and used projection-defined calcium recording and bidirectional optogenetic manipulation to determine the contribution of the S1–DMS pathway to social and cognitive deficits. We further examined three patient-associated *FMR1* missense variants expressed in S1 to determine whether distinct variants converge on shared behavioral and circuit abnormalities and whether activation of the S1–DMS pathway can ameliorate these phenotypes. Regional transcriptomic profiling of S1 and DMS was then used to identify convergent molecular alterations across *Fmr1* deficiency and the missense variants, revealing prominent changes in mitochondrial oxidative phosphorylation programs. Finally, we tested the functional relevance of mitochondrial dysfunction using two complementary approaches: systemic treatment with the mitochondria-targeted antioxidant MitoQ and spatially restricted Mitochondrial OptoEnergetic Activation (Mito-OptoEA) in S1 using mt-EcGAPR. Together, these approaches allowed us to link cross-species sensory corticostriatal abnormalities to S1– DMS circuit dysfunction, convergent mitochondrial energetic dysregulation, and *FMR1*-related social and cognitive phenotypes.

## Materials and methods

### Human participants

Participants were consecutively enrolled at the Children’s Hospital of Fudan University between January 2021 and May 2025. The final cohort comprised 52 participants with fragile X syndrome (FXS) and 64 typically developing (TD) controls. FXS was diagnosed by clinical evaluation and molecular confirmation of an *FMR1* full mutation containing more than 200 CGG repeats. Exclusion criteria were cerebral palsy, other neurological or neurodegenerative disorders, and structural lesions detected by MRI. The study was approved by the institutional ethics committee of the Children’s Hospital of Fudan University (approval no. (2021)312) and conducted in accordance with the Declaration of Helsinki. Written informed consent was obtained from each participant’s parent or legal guardian.

### Human diffusion MRI preprocessing

Diffusion MRI data and gradient tables were converted from Digital Imaging and Communications in Medicine (DICOM) to Neuroimaging Informatics Technology Initiative (NIfTI) format using dcm2niix. [39] Preprocessing was performed using MRtrix3 version 3.0.8 and FMRIB Software Library (FSL) version 6.0.7.22. [40, 41] Thermal noise was reduced using Marchenko–Pastur principal-component-analysis denoising, [42] and Gibbs-ringing artefacts were corrected by local subvoxel-shift processing. [43] Motion, eddy-current, and susceptibility-related distortions were corrected using eddy_cuda, with corresponding rotation of the b-vectors. [44] N4 bias-field correction was performed using the MRtrix3 dwibiascorrect ants workflow, [45] followed by whole-brain mask generation.

### Human tract-based spatial statistics

Diffusion tensors were fitted using an iteratively reweighted linear least-squares estimator. Fractional anisotropy (FA), mean diffusivity (MD), radial diffusivity (RD), and axial diffusivity (AD) maps were normalised to the study template and projected onto the white-matter skeleton. Voxel-wise group comparisons were performed using tract-based spatial statistics (TBSS) with threshold-free cluster enhancement (TFCE) and family-wise error (FWE) correction. [46] Age and sex were included as covariates, and TFCE/FWE-corrected *P* < 0.05 was considered statistically significant. Significant clusters were labelled using the Johns Hopkins University white-matter atlas. [47]

### Human structural-connectome analysis

Whole-brain fiber-orientation distributions were estimated using constrained spherical deconvolution, and probabilistic tractography was performed using the second-order integration over fiber orientation distributions (iFOD2) algorithm in MRtrix3. Anatomically constrained tractography incorporated tissue-type information derived from T1-weighted images. [48] One million streamlines were generated for each participant, and spherical-deconvolution informed filtering of tractograms 2 (SIFT2) was used to assign quantitative streamline weights. [49] Cortical and subcortical regions were defined using FreeSurfer and the Desikan–Killiany atlas, [50, 51] yielding 84 regions comprising 68 cortical and 16 subcortical regions. Edge weight was calculated as the sum of SIFT2 weights for streamlines connecting each region pair.

The sensorimotor–basal ganglia subnetwork comprised the bilateral precentral, postcentral, paracentral, and supramarginal cortices and the bilateral caudate, putamen, and pallidum. Of 48 possible cortical-subcortical edges, 24 contained no reconstructed streamlines across the cohort and were excluded. The remaining 24 edges were compared using two-tailed Mann– Whitney *U* tests or Welch *t*-tests, with false discovery rate (FDR) correction across edges. The displayed edge-level *P* values are uncorrected; none remained significant after FDR correction.

### Animals

*Fmr1 knockout (KO) mice* carrying the *Fmr1^tm1Cgr^*allele were obtained from The Jackson Laboratory (stock no. 004624) and backcrossed onto a C57BL/6J background for at least ten generations. Male wild-type (WT) and *Fmr1*-KO littermates were used. Behavioural experiments were conducted during the light phase. All animal procedures were approved by the Fudan University Animal Care and Use Committee (approval no. 20220329-005).

### Mouse diffusion MRI and tractography

Diffusion MRI data from seven WT and seven *Fmr1*-KO mice were analysed using an MRtrix3-based workflow. Response functions and fiber-orientation distributions were estimated from the diffusion data, and probabilistic tractography was performed using iFOD2. Pathways between S1, specified S1 subregions, and the striatum or caudoputamen (CP) were reconstructed using regions defined by the Allen Mouse Brain Common Coordinate Framework. [52] Tractography analyses were performed after excluding streamlines traversing the corpus callosum. For tractography between the entire S1 and the caudoputamen, streamlines with a minimum length of 2 mm were retained. The same procedure was applied to each of the six S1 subregion–CP pathways, using a minimum streamline length of 0.5 mm. Up to 2,000 streamlines were retained for each pathway. For each pathway, mean FA, MD, AD, and RD were calculated. Separately, a whole-brain structural connectome was constructed, and S1–CP structural connectivity strength was quantified as the sum of the SIFT2 weights assigned to the corresponding streamlines. Whole-brain mouse TBSS used TFCE inference with 1,000 permutations and a white-matter skeleton threshold of 0.25.

### Patient-associated *FMR1* missense constructs

The patient-associated human substitutions I234T, E329G, and R604Q were represented by mouse *Fmr1*-I234T, *Fmr1*-E329G, and homologous *Fmr1*-R586Q constructs, respectively. Each construct encoded a 3×FLAG epitope and a porcine teschovirus-1 2A peptide (P2A)-linked green fluorescent protein (GFP) reporter. The human synapsin (hSyn) promoter was used. Control vectors were pAAV-hSyn-GFP and pAAV-hSyn-*Fmr1*(WT)-3×FLAG-P2A-GFP. Behavioural and RNA-sequencing experiments involving I234T, E329G, and R586Q were conducted after local S1 expression in WT mice. A separate experiment in *Fmr1*-KO mice compared local S1 expression of *Fmr1*-WT and *Fmr1*-I234T.

### Stereotaxic viral injection and optical-fiber implantation

Mice were anaesthetised with isoflurane and positioned in a stereotaxic apparatus (RWD Life Science, Shenzhen, China). Viral vectors were obtained from BrainVTA, BrainCase, or Vigene Biosciences. A volume of 300 nL was delivered through a glass micropipette connected to a Nanoject III injector at 60 nL min^−1^, followed by a 10-min dwell period. Coordinates relative to bregma were anteroposterior +0.26 mm, mediolateral ±2.0 mm, and dorsoventral −1.8 mm for S1, and anteroposterior +0.26 mm, mediolateral ±2.0 mm, and dorsoventral −3.0 mm for the DMS.

Cre-dependent vectors used a double-floxed inverted open reading frame (DIO); EF1α denotes elongation factor 1α, and EYFP denotes enhanced yellow fluorescent protein. Projection-defined calcium recording used retroAAV2/R-hSyn-Cre in the DMS and AAV2/9-EF1α-DIO-GCaMP6m in S1. Axon-terminal recording used AAV2/9-hSyn-axon-GCaMP6m in S1, whereas local S1 and DMS behavioural recordings used CaMKIIα-GCaMP6m vectors driven by the calcium/calmodulin-dependent protein kinase IIα (CaMKIIα) promoter. Projection activation and inhibition used AAV2/9-hSyn-DIO-ChrimsonR-mCherry [53] and AAV2/9-EF1α-DIO-eNpHR3.0-EYFP, [54] respectively. Nonsocial S1 stimulation and recording used CaMKIIα-ChrimsonR and CaMKIIα-GCaMP6m (Supplementary Figure 4). Combined S1 activation and DMS recording used hSyn-ChrimsonR in S1 and hSyn-GCaMP6m in the DMS (Supplementary Figure 5). Missense experiments used AAV9-hSyn-GFP or the indicated AAV9-hSyn-*Fmr1*-3×FLAG-P2A-GFP construct. Mitochondrial-state perturbation used AAV9-hSyn-mt-EcGAPR-EGFP, with enhanced green fluorescent protein (EGFP) as the reporter.

### Behavioural assays

Mice were handled for approximately 2 min daily for three consecutive days and acclimated to the testing room for at least 1 h before testing. Behavioural apparatuses were cleaned with 75% ethanol between animals. Object and stranger positions were counterbalanced, and animal trajectories were analysed using Track Master version 5.0.0. The three-chamber social behaviour test was carried out as previously described. [55, 56] The three-chamber apparatus contained three interconnected chambers with 7-cm openings. Habituation, sociability, and social-novelty sessions each lasted 5 min. During sociability testing, an age-and sex-matched unfamiliar mouse was enclosed in a wire cup, and the interaction zone extended 2 cm from the cup. During social-novelty testing, the familiar and novel strangers were placed in opposite side chambers. The sociability preference index was calculated as (stranger investigation time − nonsocial-stimulus investigation time)/(stranger investigation time + nonsocial-stimulus investigation time) × 100%. The social-novelty preference index was calculated as (novel-stranger investigation time − familiar-stranger investigation time)/(novel-stranger investigation time + familiar-stranger investigation time) × 100%.

Novel-location recognition was assessed as previously described, [57] comprising five 6-min trials separated by 3-min intertrial intervals, during which mice were placed in a separate holding cage. The first trial consisted of habituation to the empty apparatus, followed by three trials with two fixed objects of identical shape, colour, and texture. In the fifth trial, one object was moved to a novel location. The recognition index was calculated as (novel-location investigation time − familiar-location investigation time)/total object-investigation time × 100%.

### Optogenetic manipulation

Optogenetic experiments were performed four weeks after bilateral viral injection and optical-fiber implantation. Yellow light was delivered bilaterally at 589 nm, 20 Hz, and 5 mW at the fiber tip. Stimulation was triggered when a mouse entered the designated social-interaction or novel-location zone. Light-off and light-on conditions were compared within the same animal. Optical output was calibrated before testing using a UNI-T UT385 photometer.

### Fiber-photometry recording and analysis

Fiber-photometry signals were acquired through ceramic-ferrule fibers with a 200-μm core using a ThinkerTech three-colour multichannel system. [58] GCaMP6m was used as the calcium indicator. [59] Signals were processed using MATLAB R2017b and expressed as Δ*F*/*F* = (F − *F*_0_)/*F*_0_ × 100%, where *F*_0_ was the pre-event baseline. Traces and heat maps were aligned to the corresponding behavioral events. Area under the curve (AUC) and peak Δ*F*/F were calculated for predefined analysis epochs. Event-level measurements were averaged within each animal before group-level analysis.

### Immunofluorescence and histology

Mice were deeply anaesthetized with isoflurane and transcardially perfused with phosphate-buffered saline followed by 4% paraformaldehyde. Brains were post-fixed overnight at 4 °C, cryoprotected in 30% sucrose, sectioned at 40 μm, and stored at −20 °C. The cryoprotectant contained 30% ethylene glycol, 30% glycerol, and 40% 0.1 M phosphate buffer at pH 7.4. Sections were blocked and permeabilized for 1 h in Tris-buffered saline containing 0.2% Triton X-100 and 3% goat serum. Primary antibody incubation was performed overnight at 4 °C, followed by fluorescent secondary antibody incubation for 1 h at room temperature in the dark.

Whole-slide images were acquired on an Olympus VS120 scanner, and confocal images were acquired on a Novel Optics NCF950 microscope using 20× or 40× objectives. GFP or FLAG fluorescence was used to confirm construct expression, and viral targeting and optical-fiber placement were assessed histologically. Animals with mistargeted expression or incorrect fiber placement were excluded from the corresponding projection analysis. Peroxisome proliferator-activated receptor gamma coactivator 1α (PGC-1α)-immunoreactive fluorescence was quantified in predefined S1 and DMS regions. Quantification included 8–9 brain sections from three mice per group, with each plotted value representing one section.

### Primary neuronal culture and Rhodamine 800 imaging

Primary hippocampal neurons were prepared from one-day-old C57BL/6 mouse pups, mechanically dissociated using fine-bore pipettes, and plated on 12-mm glass coverslips in 35-mm dishes containing 2 mL of plating medium. Per 100 mL, plating medium contained 89 mL minimum essential medium, 0.5 g glucose, 0.5 mM glutamine, 2 g sodium bicarbonate, 10 mg bovine transferrin, 2.5 mg insulin, and 10% foetal calf serum. The medium was partially replaced on day 2 and subsequently replaced twice weekly with feeding medium. Per 100 mL, feeding medium contained 97 mL minimum essential medium, 0.5 g glucose, 0.5 mM glutamine, 2 g sodium bicarbonate, 10 mg bovine transferrin, 3 μM cytarabine, and 2% B27. Neurons were transduced on day in vitro 5 with AAV-hSyn-mt-EcGAPR-EGFP or control virus.

On day in vitro 14, neurons were loaded with 50 nM Rhodamine 800 in Tyrode’s buffer for 20 min and imaged at room temperature using a ZEISS LSM 980 confocal microscope. A region encompassing one neuron was illuminated with a 532-nm laser in bleaching mode for 380 s, followed immediately by 15 min of time-series imaging. Images were processed using ImageJ version 1.54g, and background fluorescence was subtracted before quantification. The transformed Rhodamine 800 signal was calculated as −*RT*/*F* × ln (*F*_sig_/*F*_bg_), where *R* is the gas constant, *F* is the Faraday constant, *T* was 298 K, and *F*_sig_ and *F*_bg_ denote mitochondrial-region and cell-free background fluorescence, respectively. This signal was used as a relative mitochondrial membrane-potential-related recovery metric. Endpoint analysis included six control and nine mt-EcGAPR-expressing neurons. Time-course and AUC analyses included nine control and ten mt-EcGAPR-expressing neurons.

### Pharmacological mitochondria-targeted antioxidant treatment

MitoQ (1 mg/kg; Selleck Chemicals, S8978) was administered by intraperitoneal injection, and control mice received an equivalent volume of sterile saline. This dose was selected based on prior in vivo studies indicating that it is well tolerated and produces measurable mitochondrial benefits in the brain. [55, 60] Specifically, this dose has been reported to achieve blood-brain barrier penetration and to improve mitochondrial function without obvious toxicity.

### RNA-sequencing and pathway analysis

S1 and DMS tissues were collected after local expression of GFP, *Fmr1*-I234T, *Fmr1*-E329G, or *Fmr1*-R586Q. Total RNA was isolated using the FastPure Complex Tissue/Cell Total RNA Isolation Kit (Vazyme, RC113-01). Polyadenylated RNA was enriched, fragmented, reverse-transcribed, and used for library preparation. Libraries were assessed for concentration and fragment size before Illumina sequencing. Raw reads were processed using fastp version 1.3.3 for adapter removal and quality filtering.

Filtered reads were aligned to the mouse GRCm39 reference genome using HISAT2, [61] and gene-level counts were generated using featureCounts version 2.1.1 with Ensembl release 115 annotation. [62] Differential expression between each *Fmr1* missense construct and GFP control was analysed separately in S1 and DMS using DESeq2. [63] Genes with at least 10 counts in three or more libraries within each region were retained. Differentially expressed genes (DEGs) were defined by Benjamini–Hochberg-adjusted *P* < 0.05 and |log2 fold change| ≥ log2(1.5). Gene-set enrichment analysis (GSEA) was performed using DESeq2 Wald statistics with Hallmark and Gene Ontology Biological Process gene sets from Molecular Signatures Database (MSigDB; v2026.1.Mm). [64, 65] Oxidative phosphorylation (OXPHOS)-related pathways were evaluated across all six missense-versus-GFP comparisons, with FDR *q* < 0.05 considered significant. Positive normalized enrichment scores (NESs) indicated enrichment in *Fmr1* missense groups relative to GFP controls. Pathway visualization was performed using clusterProfiler. [66] Principal-component analysis, Pearson correlation analysis, DEG intersection analysis, and OXPHOS module scores before and after adjustment for cell-type-marker composition were used to evaluate transcriptomic structure (Supplementary Figure 7A–E). The RNA-sequencing experiment included eight groups (four constructs × two brain regions), with three independent bulk-tissue libraries per group.

### Statistical analysis

Data processing and statistical analyses were performed using SPSS 22, MATLAB R2017b, ImageJ 1.54g, GraphPad Prism 9.5.1 and SciPy 1.18.1. Sample sizes were selected with reference to prior mechanistic studies. [67, 68] Data distributions were assessed using Shapiro–Wilk tests. Two-group comparisons used two-tailed Student’s *t* tests, Welch *t* tests, or paired *t* tests according to the data distribution and experimental design. Multiple-group comparisons were performed using one-way or two-way analysis of variance, followed by appropriate post hoc tests including Dunnett’s, Bonferroni’s, or Tukey’s multiple-comparisons tests as specified for each experiment. Corresponding non-parametric tests were used when parametric assumptions were not met. Normality of paired differences was assessed for paired tests, and Bartlett’s test was used to assess variance homogeneity in independent-group analyses. Human and mouse TBSS used FWE correction, connectome analyses used FDR correction, and multiple-group behavioral comparisons used Dunnett’s test against GFP. Data are presented as the mean ± standard error of the mean (SEM) unless otherwise stated. The experimental unit was the participant, mouse, cultured neuron or RNA-sequencing library, as specified for each analysis. *P* < 0.05 was considered statistically significant.

### Data availability

Source data underlying the main and supplementary figure panels are available from the corresponding author upon reasonable request. RNA-sequencing data are available in the Gene Expression Omnibus under accession GSE342691. Individual-level neuroimaging data are not publicly available because of consent and privacy restrictions but may be shared in de-identified form subject to institutional approval and a data-use agreement.

## Results

### Altered sensory corticostriatal structural connectivity in patients with FXS

To characterize white-matter structural abnormalities associated with *FMR1* deficiency in the human brain, we first performed whole-brain TBSS on diffusion MRI data from 52 individuals with FXS and 64 TD controls (Figure 1A). The two groups did not differ significantly in age, whereas sex distribution differed between groups (Supplementary Figure 1). TBSS revealed widespread reductions in FA, together with extensive increases in MD and RD in FXS. These alterations prominently involved the bilateral corona radiata and internal capsule (Figure 1B, C, Supplementary Table 3, Supplementary Figure 2).

**Figure 1.**
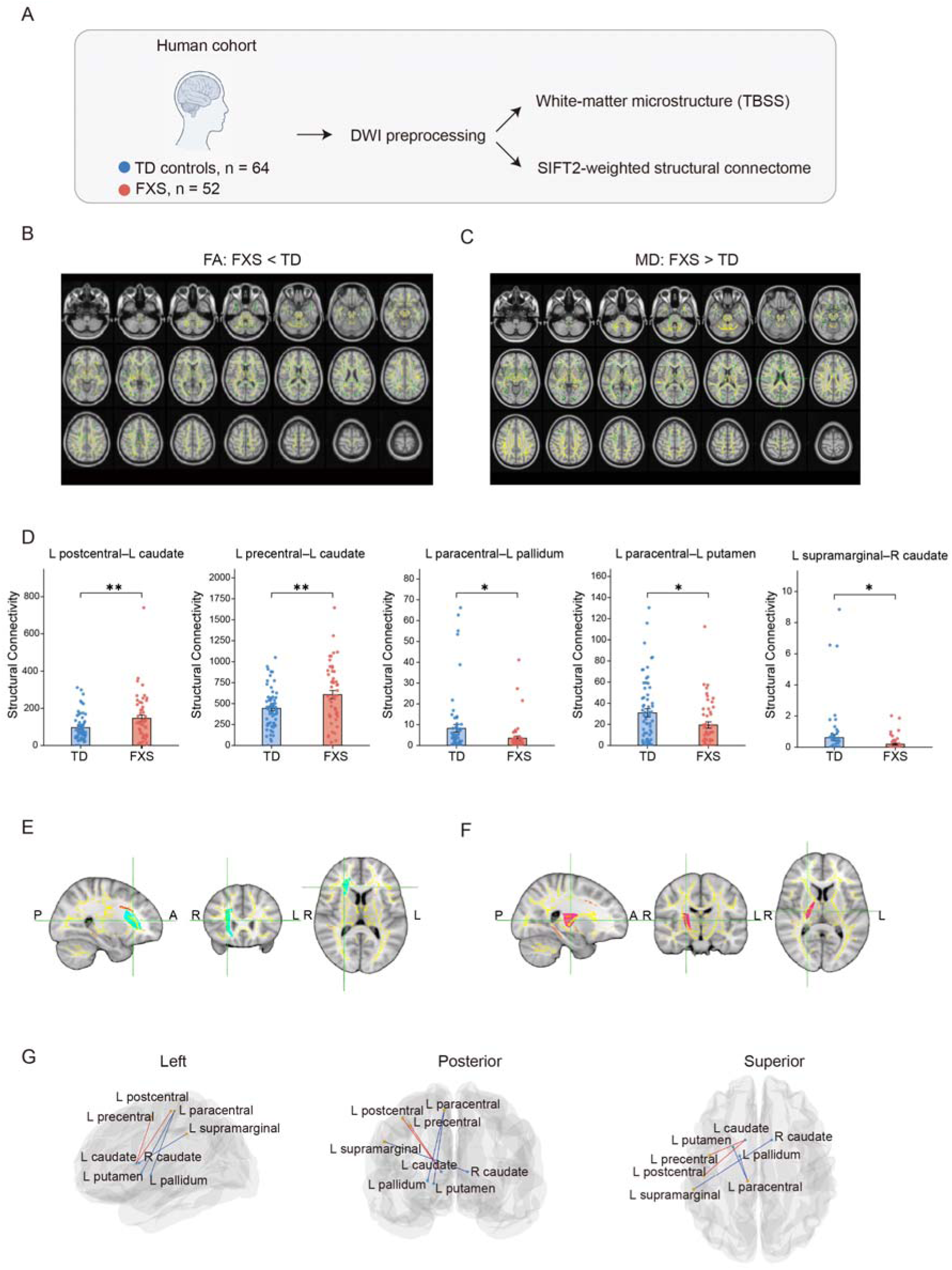
Diffusion MRI reveals white-matter and sensory corticostriatal connectivity differences in participants with FXS. (A) Human study workflow, including typically developing (TD) participants (n = 64) and participants with fragile X syndrome (FXS; n = 52), diffusion-weighted imaging (DWI) preprocessing, tract-based spatial statistics (TBSS), and SIFT2-weighted structural-connectome analysis. (B and C) TBSS maps showing lower fractional anisotropy (FA; B) and higher mean diffusivity (MD; C) in FXS than in TD participants. Significant voxels are overlaid on the white-matter skeleton and anatomical template. Analyses included age and sex as covariates and used threshold-free cluster enhancement (TFCE) with family-wise error (FWE) correction at P < 0.05. (D) SIFT2-weighted connectivity for the left postcentral–left caudate, left precentral–left caudate, left paracentral–left pallidum, left paracentral–left putamen, and left supramarginal–right caudate edges, from left to right. Symbols show individual participants; unpaired two-tailed t-tests were used for D; bars show mean ± SEM. (E and F) Anatomical overlap of reduced FA with the right corona radiata (E) and right internal capsule (F). (G) Left, posterior, and superior views of the sensorimotor–basal ganglia connectivity differences. Red and blue edges indicate higher and lower SIFT2-weighted connectivity, respectively, in FXS relative to TD participants. L, left; R, right; A, anterior; P, posterior. *P < 0.05, **P < 0.01.

Given the prominent involvement of the corona radiata and internal capsule, we next examined structural connectivity within the sensorimotor–basal ganglia network using probabilistic tractography and SIFT2-based connectome analysis. In contrast, connectivity was increased between the left caudate and the left postcentral gyrus and between the left caudate and the left precentral gyrus (Figure 1D). At the network level, connectivity was reduced between the left paracentral lobule and the left pallidum, between the left paracentral lobule and the left putamen, and between the left supramarginal gyrus and the right caudate. To visualize the anatomical distribution of these structural abnormalities, three-dimensional rendering showed the overlap of reduced FA with the right corona radiata and internal capsule (Figure 1E, F) and illustrated the pattern of sensorimotor–striatal connectivity alterations, with reduced paracentral- and supramarginal-related connections and increased primary sensorimotor–caudate connections (Figure 1G).

Together, these findings reveal altered sensory corticostriatal structural connectivity in patients with FXS, characterized predominantly by reduced connectivity across corticostriatal pathways with selective increases in primary sensorimotor–caudate connections.

### Sensory corticostriatal structural abnormalities in *Fmr1*-KO mice

To determine whether the sensory corticostriatal abnormalities identified in patients with FXS were recapitulated in a genetic mouse model, we next examined white matter organization and structural connectivity in *Fmr1*-KO mice (Figure 2A). Whole-brain TBSS revealed widespread increases in FA in *Fmr1*-KO mice, involving 12,773 voxels, whereas FA reductions were much more spatially restricted. Notably, 66.4% of the voxels showing reduced FA were localized to the internal capsule, indicating a focal alteration within a major pathway linking the cortex and striatum (Figure 2B).

**Figure 2.**
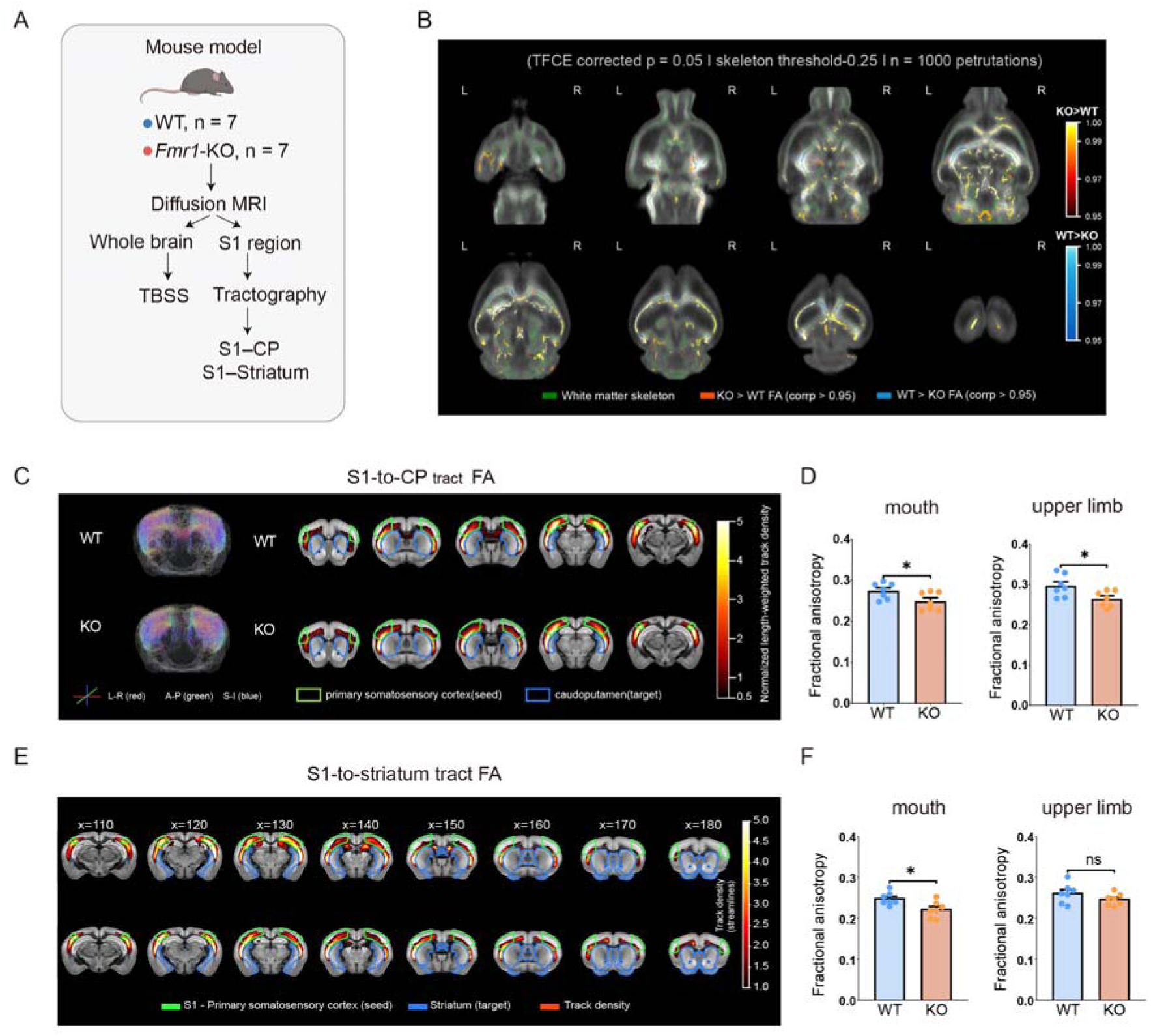
Diffusion MRI reveals sensory corticostriatal structural abnormalities in *Fmr1*-KO mice. (A) Mouse study workflow, including wild-type (WT) and *Fmr1*-knockout (KO) mice (n = 7 per genotype), whole-brain tract-based spatial statistics (TBSS), and tractography from primary somatosensory cortex (S1) to the caudoputamen (CP) and striatum. (B) Whole-brain fractional anisotropy (FA) comparison. Warm colours indicate higher FA in *Fmr1*-KO mice, and cool colours indicate higher FA in WT mice; the white-matter skeleton is shown in green. Threshold-free cluster enhancement (TFCE) used 1,000 permutations and a skeleton threshold of 0.25; corrected P < 0.05. (C) Representative S1-to-CP tractography and tract-density maps in WT and *Fmr1*-KO mice. The S1 seed and CP target are outlined in green and blue, respectively; the colour scale indicates normalised length-weighted track density. (D) FA in the mouth and upper-limb S1-to-CP pathways. (E) Representative S1-to-striatum tract-density maps, with the S1 seed in green and striatal target in blue. (F) FA in the mouth and upper-limb S1-to-striatum pathways. Individual points represent mice; n = 7 per genotype in D and F. Unpaired two-tailed t-tests were used for D and F. Data are presented as mean ± SEM. *P < 0.05; ns, not significant.

We therefore examined projections from the S1 to the CP in greater detail (Figure 2C). FA was reduced across the overall S1–CP projection, with significant decreases in the mouth and upper-limb S1–CP projections (Figure 2C, D, Supplementary Figure 3A). Reduced FA was similarly observed in the mouth S1–striatal projection with no difference in upper-limb (Figure 2E, F, Supplementary Figure 3B), demonstrating that microstructural abnormalities extended across multiple somatotopically defined sensory corticostriatal pathways.

Thus, sensory corticostriatal pathways in *Fmr1*-KO mice showed reduced tract FA across multiple S1 projections together with increased structural connectivity in selected connections. These findings reveal altered sensory corticostriatal structural connectivity in *Fmr1*-KO mice and parallel the pathway-level abnormalities observed in patients with FXS.

### *FMR1* deficiency disrupts state-dependent recruitment of the S1– DMS pathway

Given the structural alterations detected in S1–striatal pathways, we next examined whether *FMR1* deficiency affects the functional recruitment of the S1–DMS pathway across distinct behavioral states. We first assessed spatial recognition memory using the novel-location recognition test. Consistent with previous reports of learning deficits induced by *Fmr1* deficiency, [35, 69] *Fmr1*-KO mice showed impaired novel-location recognition relative to WT littermates, with reduced exploration of the object placed in a novel location (Figure 3A–C). To determine how S1 and striatal neurons are recruited during exploration of a novel object, we expressed AAV-CaMKIIα-GCaMP6m in S1 and recorded calcium activity from local excitatory neurons. Object-evoked S1 calcium responses were comparable between WT and *Fmr1*-KO mice (Figure 3D). By contrast, local DMS recordings revealed increased calcium responses in *Fmr1*-KO mice during object contact (Figure 3E). We next examined projection-defined S1 neurons by injecting retro-AAV-hSyn-Cre into the DMS together with AAV-EF1α-DIO-GCaMP6m into S1. Calcium responses in DMS-projecting S1 neurons showed a trend toward an increase in *Fmr1*-KO mice during object exploration (Figure 3F). Consistently, when axon-targeted GCaMP6m was expressed in S1 and terminal-associated signals were recorded in the DMS, object-evoked responses were significantly increased in *Fmr1*-KO mice (Figure 3G). Thus, *FMR1* deficiency did not produce a generalized increase in local S1 recruitment during object exploration, but was associated with enhanced downstream engagement of the S1–DMS pathway and DMS.

**Figure 3.**
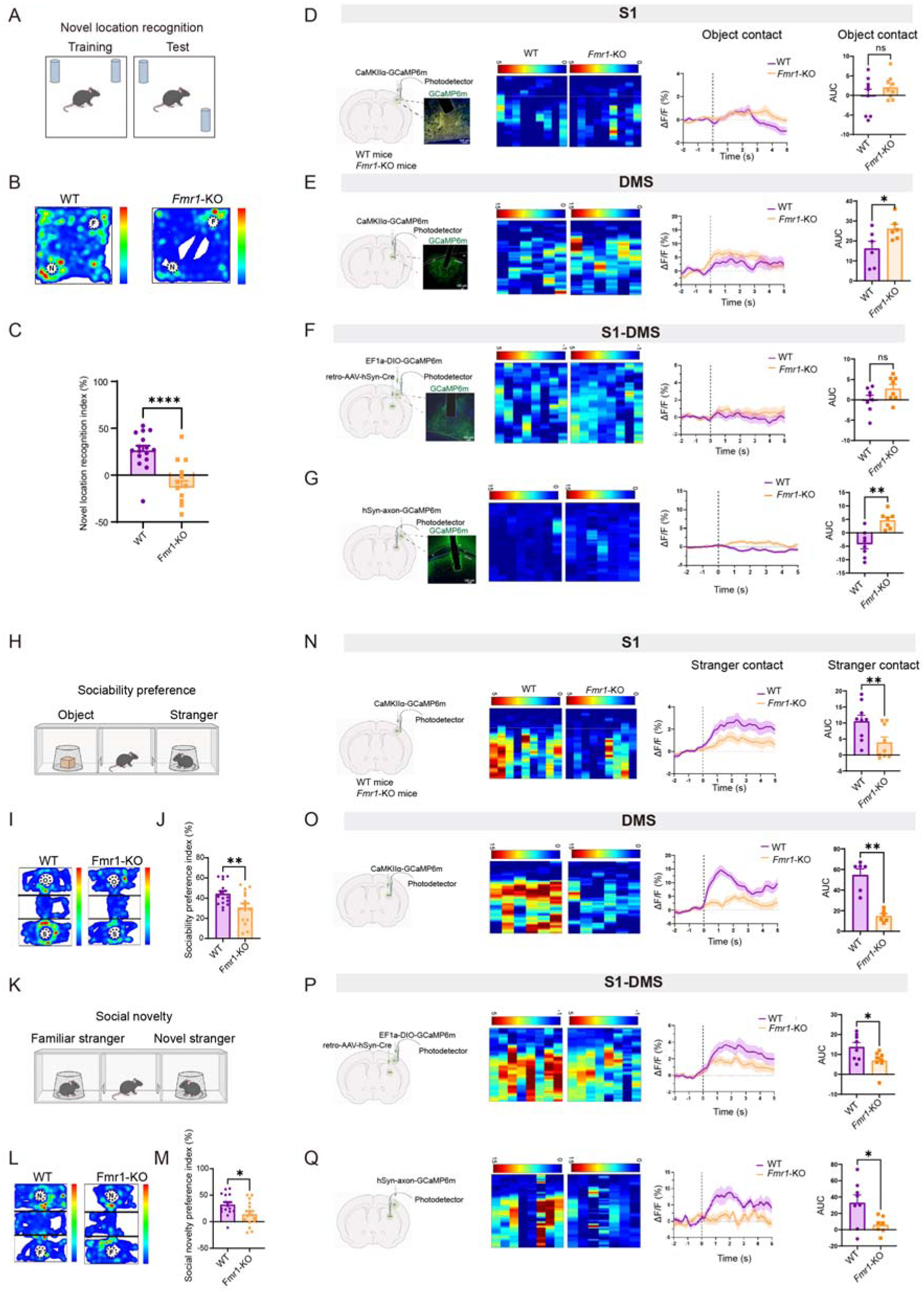
*FMR1* deficiency disrupts behavioural performance and state-dependent recruitment of the S1–DMS pathway. (A–C) Novel-location-recognition task schematic (A), representative occupancy heat maps (B), and recognition indices (C; WT, n = 16 mice; *Fmr1*-KO, n = 14 mice). (D–G) Recording configurations, representative GCaMP6m expression, animal-level heat maps, peri-event ΔF/F traces, and area under the curve (AUC) aligned to object contact. Recordings were obtained from local S1 neurons expressing CaMKIIα-GCaMP6m (D; n = 9 mice per genotype), local DMS neurons expressing CaMKIIα-GCaMP6m (E; n = 6 mice per genotype), DMS-projecting S1 neurons labelled using retrograde Cre delivery to DMS and Cre-dependent GCaMP6m expression in S1 (F; n = 8 mice per genotype), and S1 axon terminals in DMS labelled using hSyn-axon-GCaMP6m (G; WT, n = 8 mice; *Fmr1*-KO, n = 7 mice). (H–J) Three-chamber sociability task schematic (H), representative occupancy heat maps (I), and sociability preference indices (J; WT, n = 16 mice; *Fmr1*-KO, n = 15 mice). (K– M) Social-novelty task schematic (K), representative occupancy heat maps (L), and social-novelty preference indices (M; WT, n = 16 mice; *Fmr1*-KO, n = 15 mice). (N–Q) Recording configurations, animal-level heat maps, peri-event ΔF/F traces, and AUC aligned to stranger contact for local S1 neurons (N; n = 9 mice per genotype), local DMS neurons (O; n = 6 mice per genotype), DMS-projecting S1 neurons (P; n = 8 mice per genotype), and S1 axon terminals in DMS (Q; WT, n = 8 mice; *Fmr1*-KO, n = 7 mice). Unpaired two-tailed t-tests were used for C–G, J, M, and P; two-tailed Mann–Whitney U tests were used for N and O; Welch’s unpaired two-tailed t-test was used for Q. Scale bars, 100 μm. Data are presented as mean ± SEM. *P < 0.05, **P < 0.01, ****P < 0.0001; ns, not significant. N and F in the behavioural heat maps denote novel and familiar locations or strangers, respectively; O and S denote object and stranger.

We next examined S1–DMS recruitment during social behavior. In the three-chamber test, *Fmr1*-KO mice showed reduced preference for an unfamiliar conspecific over the nonsocial stimulus (Figure 3H–J) and reduced preference for a novel over a familiar conspecific (Figure 3K–M), consistent with previous reports. [35] In contrast to the pattern observed during object exploration, social interaction produced a coordinated reduction in neuronal recruitment across the S1–DMS pathway. Local S1 and DMS calcium responses to stranger contact were both markedly reduced in *Fmr1*-KO mice relative to WT controls (Figure 3N,O). This reduction was more evident in DMS-projecting S1 neurons and in S1 terminal-associated calcium signals recorded in the DMS (Figure 3P,Q), indicating that impaired social recruitment was expressed consistently at both the local and projection levels of the S1–DMS pathway. Together, these findings show that *FMR1* deficiency disrupts the state-dependent recruitment of the S1–DMS pathway, with a particularly pronounced reduction during social interaction.

### S1–DMS projection activation improves behavioral abnormalities across *FMR1*-deficient and variant models

To determine whether the S1–DMS projection causally regulates social behavior, we used a retrograde Cre-dependent strategy to manipulate DMS-projecting S1 neurons during the three-chamber sociability test. In *Fmr1*-KO mice expressing ChrimsonR in the S1–DMS projection, optogenetic activation significantly increased sociability preference relative to the laser-off condition (Figure 4A–D), whereas inhibition of the same projection in WT mice reduced sociability preference (Figure 4E–H).

**Figure 4.**
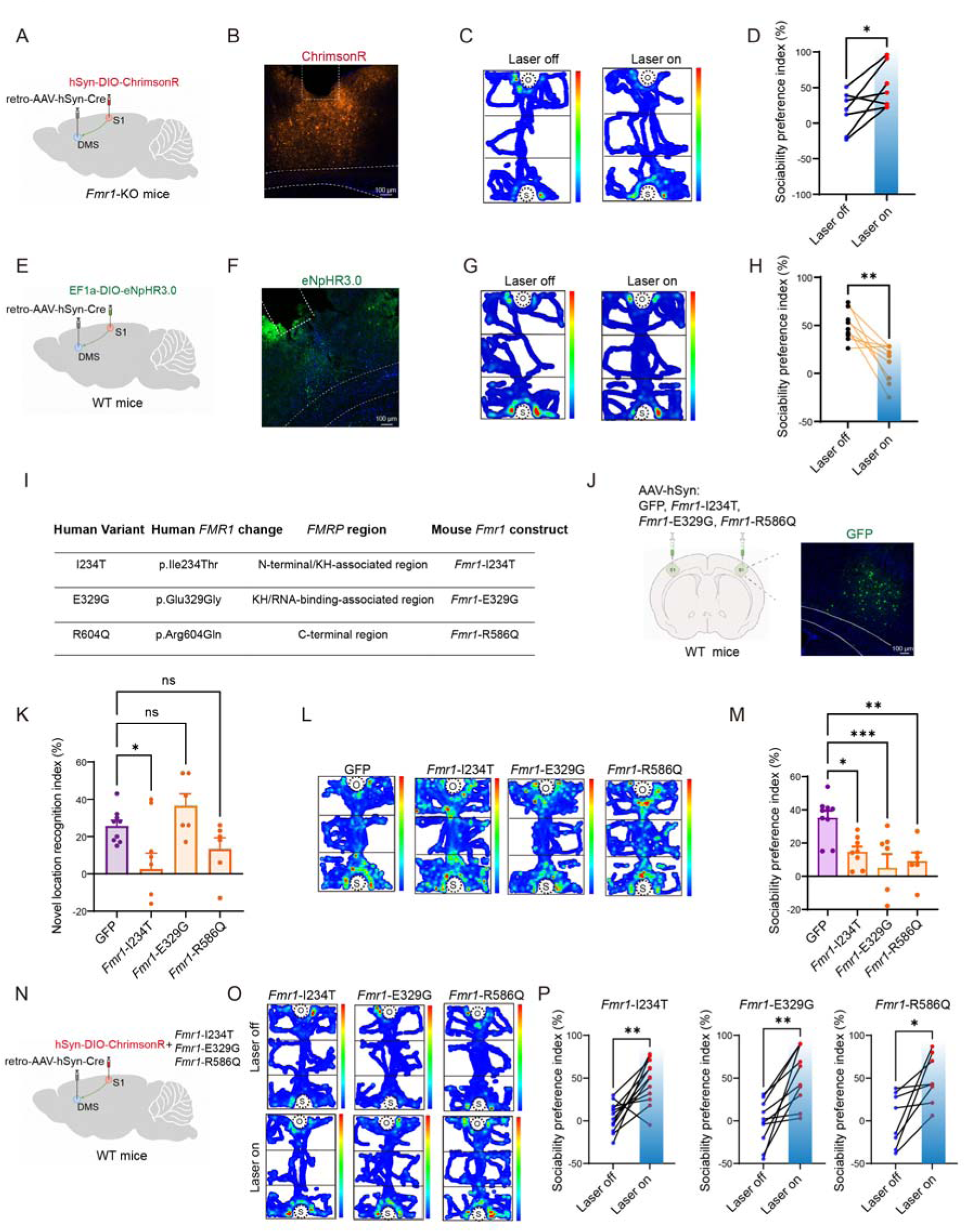
S1–DMS projection manipulation and S1-targeted *Fmr1* missense variants alter behavioural performance. (A–D) S1–DMS projection activation in *Fmr1*-KO mice: viral and optical configuration using retrograde Cre delivery to DMS and Cre-dependent ChrimsonR expression in S1 (A), representative ChrimsonR expression (B), occupancy heat maps during laser-off and laser-on testing (C), and paired sociability preference indices (D; n = 7 mice). (E–H) S1–DMS projection inhibition in WT mice: retrograde Cre and Cre-dependent eNpHR3.0 configuration (E), representative eNpHR3.0 expression (F), occupancy heat maps (G), and paired sociability preference indices (H; n = 9 mice). (I) Patient-associated human *FMR1* substitutions I234T, E329G, and R604Q, the corresponding FMRP regions, and mouse *Fmr1* constructs encoding I234T, E329G, and the homologous R586Q substitution. (J) S1 expression of GFP or the indicated *Fmr1* construct in WT mice, with representative GFP fluorescence. (K) Novel-location-recognition indices (GFP, n = 9 mice; *Fmr1*-I234T, n = 9 mice; *Fmr1*-E329G, n = 6 mice; *Fmr1*-R586Q, n = 6 mice). (L and M) Representative sociability occupancy heat maps (L) and preference indices (M; GFP, n = 9 mice; *Fmr1*-I234T, n = 8 mice; *Fmr1*-E329G, n = 7 mice; *Fmr1*-R586Q, n = 6 mice). (N–P) S1–DMS projection activation in WT mice expressing *Fmr1*-I234T, *Fmr1*-E329G, or *Fmr1*-R586Q in S1: viral and optical configuration (N), occupancy heat maps during laser-off and laser-on testing (O), and paired sociability preference indices (P; *Fmr1*-I234T, n = 12 mice; *Fmr1*-E329G, n = 10 mice; *Fmr1*-R586Q, n = 8 mice). Two-tailed paired t-tests were used for D, H, and P. One-way analysis of variance followed by Dunnett’s multiple-comparisons test against GFP was used for K and M. Scale bars, 100 μm. Data are presented as mean ± SEM. *P < 0.05, **P < 0.01; ns, not significant. O, object; S, stranger.

We next examined whether patient-associated *FMR1* variants produced similar behavioral abnormalities. We generated three mouse *Fmr1* point-mutant constructs corresponding to the human p.Ile234Thr, p.Glu329Gly and p.Arg604Gln variants (*Fmr1*-I234T, *Fmr1*-E329G and *Fmr1*-R586Q, respectively) and expressed each construct locally in S1 neurons of WT mice (Figure 4I,J). Mice expressing *Fmr1*-I234T showed impaired novel-location recognition, whereas mice expressing *Fmr1*-E329G or *Fmr1*-R586Q did not exhibit significant deficits (Figure 4K), indicating variant-dependent effects on spatial recognition. Consistently, in *Fmr1*-KO mice, S1-targeted expression of *Fmr1*-I234T resulted in a significantly lower novel-location recognition index than expression of *Fmr1*-WT (Supplementary Figure 6A,B). By contrast, all three variants significantly reduced sociability preference relative to GFP controls (Figure 4L,M), recapitulating the social abnormality observed in *Fmr1*-KO mice (Figure 3H–J). In *Fmr1*-KO mice, S1-targeted expression of *Fmr1*-I234T also resulted in a significantly lower social recognition index than expression of *Fmr1*-WT (Supplementary Figure 6C,D). Thus, patient-associated *FMR1* variants converged on impaired sociability while producing distinct effects on spatial recognition, with the I234T variant affecting both behavioral domains.

We then asked whether increasing S1–DMS activity could improve these variant-associated behavioral abnormalities. S1–DMS activation significantly increased sociability preference in mice expressing *Fmr1*-I234T, *Fmr1*-E329G or *Fmr1*-R586Q in S1 (Figure 4N–P). To further test whether this effect extended to spatial recognition, we co-expressed *Fmr1*-I234T and AAV-hSyn-DIO-ChrimsonR in S1 together with retro-AAV-hSyn-Cre in the DMS, enabling projection-specific activation of S1–DMS neurons in the I234T model. During novel-location exploration, S1–DMS activation significantly increased the recognition index relative to the laser-off condition (Supplementary Figure 6E,F).

Together, these findings show that S1–DMS activity bidirectionally regulates sociability and that enhancing this pathway improves behavioral abnormalities across *FMR1*-deficient and missense-variant models.

### Patient-associated *FMR1* variants converge on oxidative phosphorylation-related transcriptional programs in S1 and DMS

To investigate molecular mechanisms associated with the behavioral abnormalities induced by patient-associated *FMR1* variants, we performed bulk RNA sequencing of S1 and DMS tissue following local S1 expression of GFP, *Fmr1*-I234T, *Fmr1*-E329G or *Fmr1*-R586Q (Figure 5A). Differential gene expression varied substantially among variants and between regions. In S1, I234T, E329G and R586Q resulted in 54, 30 and 6 upregulated DEGs and 157, 353 and 90 downregulated DEGs, respectively. In DMS, the corresponding comparisons identified 161, 12 and 11 upregulated DEGs and 213, 71 and 90 downregulated DEGs (Figure 5B).

**Figure 5.**
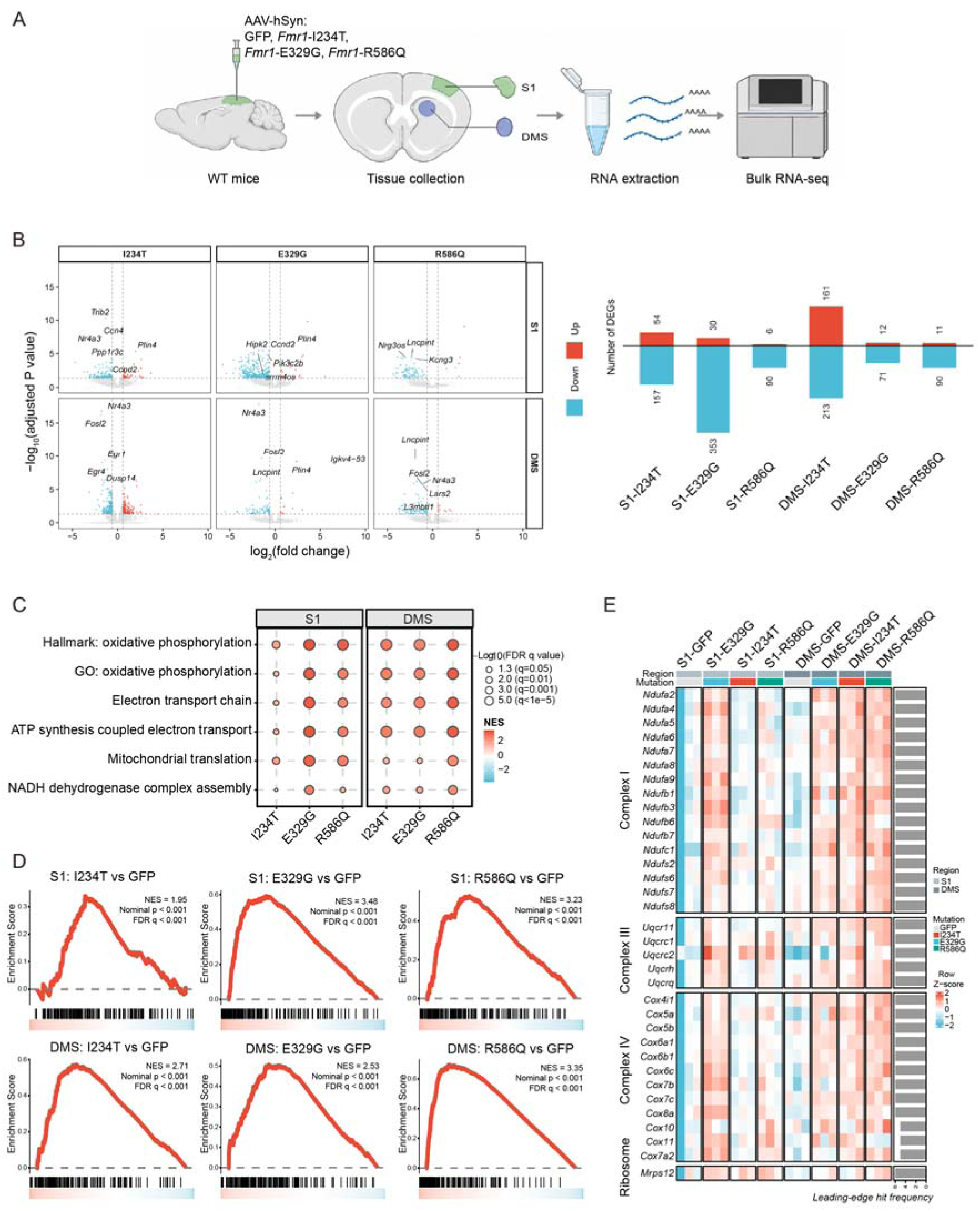
Bulk RNA sequencing identifies region- and construct-specific differential expression and shared OXPHOS pathway enrichment. (A) Experimental workflow. GFP or the indicated *Fmr1* construct (*Fmr1*-I234T, *Fmr1*-E329G, or *Fmr1*-R586Q) was expressed locally in S1 of WT mice; S1 and DMS tissues were then collected for bulk RNA sequencing. (B) Volcano plots and numbers of upregulated and downregulated differentially expressed genes (DEGs). Counts were 54 and 157 for I234T, 30 and 353 for E329G, and 6 and 90 for R586Q in S1; the corresponding counts in DMS were 161 and 213, 12 and 71, and 11 and 90. (C) Oxidative phosphorylation (OXPHOS)-related pathway enrichment in S1 and DMS. Bubble size indicates FDR-adjusted significance, and bubble colour indicates the normalised enrichment score (NES). (D) Hallmark OXPHOS gene-set enrichment analysis (GSEA) curves for six construct-versus-GFP contrasts across S1 and DMS. NES values for I234T, E329G, and R586Q were 1.95, 3.48, and 3.23, respectively, in S1 and 2.71, 2.53, and 3.35, respectively, in DMS. (E) Heat map of genes associated with respiratory-chain complexes I, III, and IV and mitochondrial ribosomes. Colour indicates the row Z-score; the adjoining bars show leading-edge hit frequency. Each region-by-construct group comprised n = 3 independent bulk-tissue libraries. DEGs were identified using DESeq2, and gene-set enrichment was analysed using GSEA. GO, Gene Ontology.

Despite these differences in DEG number and expression pattern, GSEA consistently identified positive enrichment of OXPHOS-related pathways across all six region-specific variant comparisons. Enriched gene sets included Hallmark oxidative phosphorylation, oxidative phosphorylation, electron transport chain, ATP synthesis-coupled electron transport, mitochondrial translation and reduced nicotinamide adenine dinucleotide (NADH) dehydrogenase complex assembly, although the specific pathways and enrichment magnitude varied among variants (Figure 5C). Notably, the Hallmark OXPHOS gene set was positively enriched for all three variants in both S1 and DMS (Figure 5D).

A heat map of selected OXPHOS-related genes further revealed coordinated alterations involving respiratory-chain complexes I, III and IV and mitochondrial ribosomal genes across the region-by-variant groups (Figure 5E). The OXPHOS module-score pattern remained evident after adjustment for cell-type-marker composition (Supplementary Figure 7E).

Together, these findings indicate that distinct patient-associated *FMR1* variants converge on OXPHOS-related transcriptional remodeling in both S1 and DMS despite substantial variant- and region-dependent transcriptional heterogeneity.

### Pharmacological and S1 Mito-OptoEA targeting of mitochondrial energetic state improves *FMR1*-related behavioral abnormalities

Given the recurrent enrichment of OXPHOS-related transcriptional programs, we next asked whether targeting mitochondrial energetic state could modify *FMR1*-related behavioral abnormalities. We first treated *Fmr1*-KO mice and their WT littermates with MitoQ, a mitochondria-targeted antioxidant (i.p., 1 mg/kg), daily for 7 days, with MitoQ administration continued throughout behavioral testing (Figure 6A). In the novel location test, vehicle-treated *Fmr1*-KO mice showed impaired novel location recognition, whereas MitoQ-treated *Fmr1*-KO mice exhibited a significantly greater preference for the object placed in the novel location (Figure 6B). In the three-chamber social interaction test, vehicle-treated *Fmr1*-KO mice showed reduced social interaction, whereas MitoQ treatment significantly increased their interaction with the social stimulus (Figure 6C). In the subsequent social novelty test, vehicle-treated *Fmr1*-KO mice displayed reduced preference for the novel social stimulus, whereas MitoQ-treated *Fmr1*-KO mice showed a significantly greater preference for the novel mouse (Figure 6D). Thus, systemic pharmacological targeting of mitochondrial function attenuated deficits in novel location recognition, social interaction, and social novelty in *Fmr1*-KO mice.

**Figure 6.**
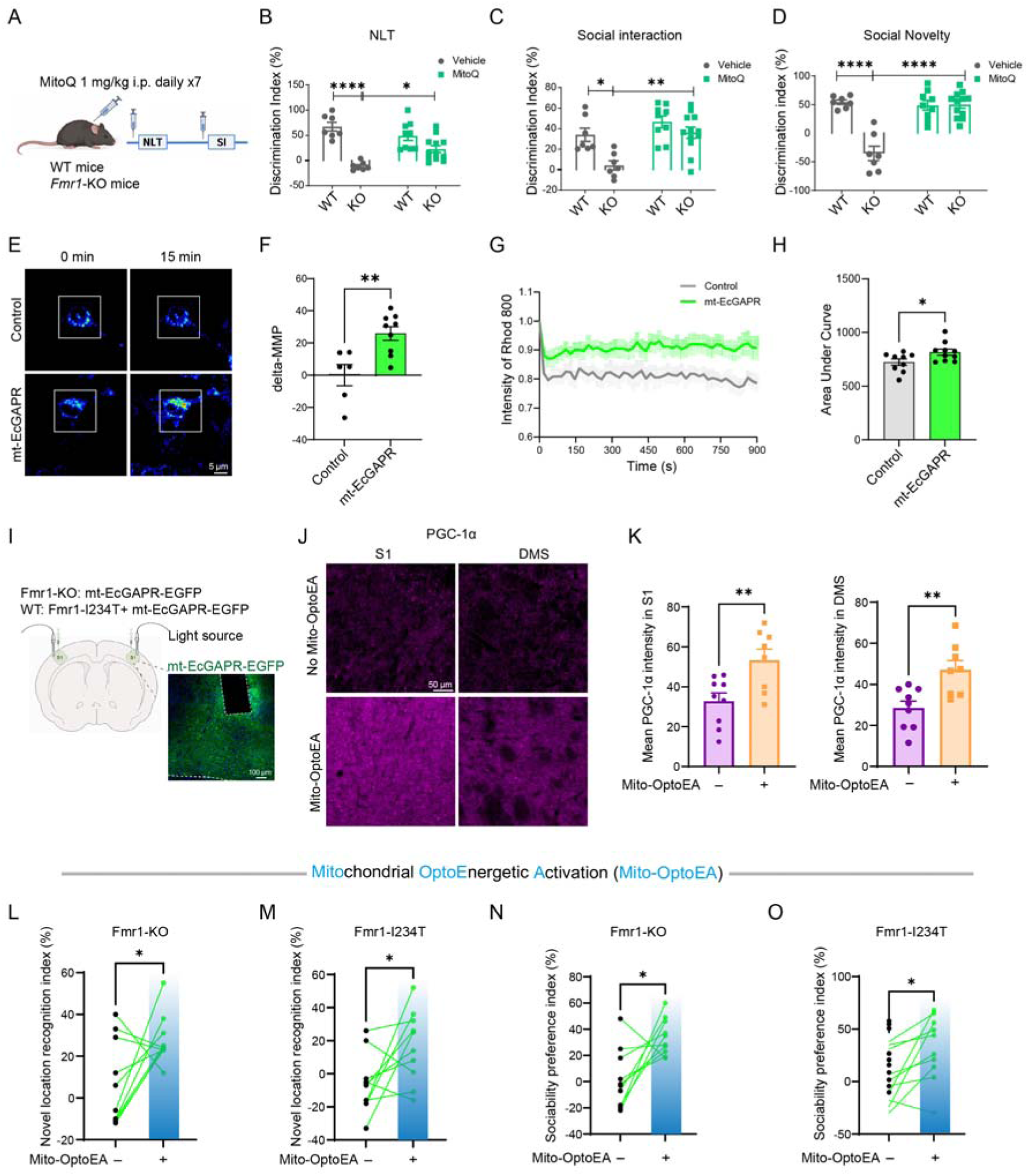
Pharmacological and S1 Mito-OptoEA targeting of mitochondrial function attenuates *FMR1*-related behavioral abnormalities. (A) Experimental timeline for MitoQ administration (1 mg/kg, intraperitoneally, daily for 7 days) before behavioural testing, with treatment continued during testing. NLT, novel-location test; SI, social interaction. (B) Discrimination indices in the novel-location test. (C) Discrimination indices in the three-chamber social-interaction test. (D) Discrimination indices in the social-novelty test. In B–D, wild-type (WT) and *Fmr1*-knockout (KO) mice received vehicle or MitoQ (vehicle: WT, n = 7; *Fmr1*-KO, n = 7; MitoQ: WT, n = 9; *Fmr1*-KO, n = 12). (E) Representative Rhodamine 800 pseudocolour images acquired immediately (0 min) and 15 min after illumination in control and mt-EcGAPR-expressing primary hippocampal neurons; boxes indicate the illuminated regions (scale bar, 5 μm). (F) Endpoint change in the transformed Rhodamine 800 signal (control, n = 6 neurons; mt-EcGAPR, n = 9 neurons). (G) Normalised Rhodamine 800 fluorescence during the 900-s recovery period (control, n = 9 neurons; mt-EcGAPR, n = 10 neurons). (H) AUC of the normalised Rhodamine 800 recovery signal (control, n = 9 neurons; mt-EcGAPR, n = 10 neurons). (I) S1-targeted mt-EcGAPR expression and illumination in *Fmr1*-KO mice and WT mice with local S1 expression of *Fmr1*-I234T, with representative mt-EcGAPR-EGFP fluorescence (scale bar, 100 μm). (J) Representative PGC-1α immunofluorescence in S1 and DMS with and without Mitochondrial OptoEnergetic Activation (Mito-OptoEA) in the *Fmr1*-I234T model (scale bar, 50 μm). (K) Mean PGC-1α immunofluorescence intensity per section in S1 and DMS (without Mito-OptoEA, n = 9 sections; with Mito-OptoEA, n = 8 sections; three mice per group). (L and M) Paired novel-location-recognition indices in *Fmr1*-KO mice (L) and *Fmr1*-I234T-expressing WT mice (M). (N and O) Paired sociability preference indices in *Fmr1*-KO mice (N) and *Fmr1*-I234T-expressing WT mice (O). Behavioural comparisons used light-off and light-on conditions, with n = 10 mice per experiment. Two-way ANOVA followed by Tukey’s multiple comparisons test were used for B, C, and D; Unpaired two-tailed t-tests were used for F, H, and K; two-tailed paired t-tests were used for L–O. The Rhodamine 800 signal was used as a relative recovery readout associated with mitochondrial membrane potential (MMP). *P < 0.05, **P < 0.01, ****P < 0.0001. Data in B-D, F–H and K–O are presented as mean ± SEM.

We next sought to determine whether a spatially restricted and temporally controlled enhancement of neuronal mitochondrial function in S1 could similarly modify *FMR1*-related phenotypes. Based on our previous study, [37] we generated an hSyn-driven, neuron-specific mt-EcGAPR construct, packaged it into AAV, and first validated its effects in primary neurons. Following 532-nm green-light illumination, mt-EcGAPR-expressing neurons showed a greater change in Rhodamine 800 fluorescence at the 15-min endpoint than control-virus neurons (Figure 6E, F). Rhodamine 800 fluorescence recovery over the subsequent 900 s was also increased, as reflected by a greater recovery AUC (Figure 6G, H). These results indicate that green-light activation of mt-EcGAPR increased mitochondrial membrane potential in neurons. We expressed mt-EcGAPR in S1 of mice with local S1 expression of *Fmr1*-I234T and applied green-light stimulation, thereby establishing an in vivo S1 Mito-OptoEA paradigm (Figure 6I). To determine whether this manipulation altered mitochondrial signaling within the targeted region and its downstream corticostriatal target, we measured PGC-1α expression in S1 and DMS. PGC-1α immunofluorescence intensity was significantly increased in S1 following Mito-OptoEA compared with the corresponding control condition. Notably, PGC-1α immunofluorescence was also increased in the downstream DMS (Figure 6J, K), indicating that S1 Mito-OptoEA enhanced mitochondrial biogenesis-related signaling both locally in S1 and in a downstream corticostriatal region.

We then tested whether S1 Mito-OptoEA altered *FMR1*-related behavioral abnormalities. Green-light activation of mt-EcGAPR significantly increased the novel-location recognition index in both *Fmr1*-KO mice and mice with S1 expression of *Fmr1*-I234T compared with the corresponding light-off conditions (Figure 6L, M). In the three-chamber social interaction test, Mito-OptoEA similarly increased sociability preference in both *Fmr1*-KO mice and mice expressing *Fmr1*-I234T relative to the corresponding light-off conditions (Figure 6N, O).

Together, these findings show that both systemic pharmacological intervention with MitoQ and spatially restricted S1 Mito-OptoEA targeting of mitochondrial energetic state attenuated *FMR1*-related social and cognitive behavioral abnormalities.

## Discussion

Our study establishes a sensory corticostriatal circuit–metabolic framework linking *FMR1* perturbation to social and cognitive dysfunction in FXS. Cross-species diffusion MRI identified disrupted sensory corticostriatal structural connectivity in individuals with FXS and *Fmr1*-KO mice, with mouse analyses further localizing abnormalities to S1–striatal pathways. Functional interrogation then demonstrated that dysfunction of the S1–DMS pathway contributes to social and cognitive deficits. Three patient-associated *FMR1* variants converged on social behavioral deficits, which were improved by S1–DMS activation. Transcriptomic analyses further identified convergent OXPHOS-related mitochondrial remodeling in S1 and DMS across these variants, and Mito-OptoEA improved social and cognitive deficits in FXS mouse models. Together, these findings connect cross-species structural abnormalities to a defined corticostriatal circuit and a convergent mitochondrial mechanism.

Cross-species neuroimaging provides a means to connect clinically detectable brain structural abnormalities with experimentally accessible circuits in genetic models of neurodevelopmental disorders. [70] In individuals with FXS, we identified widespread white-matter abnormalities with prominent involvement of sensorimotor–basal ganglia pathways. Diffusion MRI in *Fmr1*-KO mice further localized structural abnormalities to S1–striatal pathways, providing a cross-species anatomical basis for subsequent interrogation of the S1– DMS projection. These findings extend prior evidence of white-matter and structural-network abnormalities in FXS by linking the systems-level phenotype to a sensory corticostriatal pathway that can be directly examined in the mouse model.

At the circuit level, our findings identify the S1–DMS projection as a functional substrate of *FMR1*-associated social and cognitive deficits. Corticostriatal abnormalities have been reported in neurodevelopmental-disorder models, including early corticostriatal hyperactivity in *Shank3B*-deficient mice and cell-type-specific disruption of corticostriatal circuitry in *Fmr1*-KO mice. [71, 72] Here, *Fmr1*-KO mice showed reduced recruitment of DMS-projecting S1 neurons and their terminals in DMS during social interaction. Bidirectional manipulation further demonstrated a causal contribution of S1–DMS output to social behavior, while activation of this projection improved sociability in *Fmr1*-KO mice and across all three patient-associated *FMR1* variant models and improved novel-location recognition in the I234T model. The DMS contains heterogeneous populations of spiny projection neurons and interneurons with distinct input–output relationships and behavioral functions, [73, 74] and striatal photometry can include substantial nonsomatic calcium signals. [75] Resolving this circuit architecture will require cell-type-specific recordings and manipulations together with direct analysis of S1–DMS synaptic connectivity.

Our mitochondrial findings identify a convergent cellular mechanism within this sensory corticostriatal framework. Previous studies have linked FMRP dysfunction to abnormal mitochondrial biogenesis, energetic regulation, ATP-synthase proton leak, and altered mitochondrial dynamics. [33, 36–38] Across all three patient-associated *FMR1* variants, transcriptomic analyses revealed recurrent enrichment of OXPHOS-related programs in both S1 and DMS, indicating convergent mitochondrial remodeling across genetically distinct *FMR1* perturbations. We further developed an mt-EcGAPR-based Mito-OptoEA system to directly manipulate neuronal mitochondrial energetic state in S1. Green-light activation of mt-EcGAPR increased mitochondrial membrane potential in primary neurons, while S1-targeted Mito-OptoEA improved sociability and novel-location recognition in *Fmr1*-KO and I234T models. The accompanying increase in PGC-1α in both S1 and DMS further supports a mitochondrial response within the sensory corticostriatal system. These findings identify mitochondrial energetic state as a modifiable component of *FMR1*-associated behavioral dysfunction.

Together, our study defines a cross-species sensory corticostriatal circuit–metabolic framework for FXS, linking S1–DMS circuit dysfunction and convergent mitochondrial energetic remodeling to social and cognitive deficits. Targeting circuit activity and mitochondrial energetics may therefore provide tractable strategies for intervention.

## Author contributions

MS and MS conceived and designed the study. YY, DL, DF, JY, MS, PL, FZ, JC, and BY performed the experiments and collected the data. MS and YY prepared the visualizations. MS, RL, QX, and PL acquired funding. MS, RL, YH, QX, and YJ supervised the research. MS drafted the manuscript. MS, YY, and MS reviewed and edited the manuscript. The order of the co–first authors was determined by their relative contributions to the study, as agreed upon by all co–first authors.

## Acknowledgements

We thank Dr. Fei Li at Xinhua Hospital, Shanghai Jiao Tong University School of Medicine, Dr. Jian-Sheng Kang at the First Affiliated Hospital of Zhengzhou University, Dr. Xiaobing Yuan and Dr. Lujian Liao at East China Normal University, Dr. Jiayi Zhang, Dr. Ying Liu, Dr. Huijuan Wu and Dr. Weiguang Li at Fudan University for invaluable support and for providing the mouse lines and experimental tools; Dr. Jianzhi Wang, Junying Liu and Fan Wang at the School of Basic Medical Sciences, Dr. Min Jiang, Ying Shi and Qian Huang at the MOE Frontiers Center for Brain Science for core facility and technical support; Dr. Man Xiong and Dr. Feifei Wang for input on the manuscript.

## Supplementary material

Supporting Information includes key resources (Supplementary Table 1) and statistical analyses (Supplementary Table 2) and is available from the Wiley Online Library or from the author.

## References

1. R. J. Hagerman, E. Berry-Kravis, H. C. Hazlett, et al., “Fragile X syndrome,” Nature Reviews Disease Primers 3, no. 1 (2017): 17065. 10.1038/nrdp.2017.65.

2. A. J. Verkerk, M. Pieretti, J. S. Sutcliffe, et al., “Identification of a gene (FMR-1) containing a CGG repeat coincident with a breakpoint cluster region exhibiting length variation in fragile X syndrome,” Cell 65, no. 5 (1991): 905–914. 10.1016/0092-8674(91)90397-h.

3. J. C. Darnell, S. J. Van Driesche, C. Zhang, et al., “FMRP stalls ribosomal translocation on mRNAs linked to synaptic function and autism,” Cell 146, no. 2 (2011): 247–261. 10.1016/j.cell.2011.06.013.

4. M. Ascano, Jr., N. Mukherjee, P. Bandaru, et al., “FMRP targets distinct mRNA sequence elements to regulate protein expression,” Nature 492, no. 7429 (2012): 382–386. 10.1038/nature11737.

5. J. D. Richter and X. Zhao, “The molecular biology of FMRP: new insights into fragile X syndrome,” Nature Reviews Neuroscience 22, no. 4 (2021): 209–222. 10.1038/s41583-021-00432-0.

6. K. M. Huber, S. M. Gallagher, S. T. Warren, and M. F. Bear, “Altered synaptic plasticity in a mouse model of fragile X mental retardation,” Proceedings of the National Academy of Sciences 99, no. 11 (2002): 7746–7750. 10.1073/pnas.122205699.

7. G. Dölen, E. Osterweil, B. S. Rao, et al., “Correction of fragile X syndrome in mice,” Neuron 56, no. 6 (2007): 955–962. 10.1016/j.neuron.2007.12.001.

8. J. T. Gonçalves, J. E. Anstey, P. Golshani, and C. Portera-Cailliau, “Circuit level defects in the developing neocortex of Fragile X mice,” Nature Neuroscience 16, no. 7 (2013): 903– 909. 10.1038/nn.3415.

9. E. G. Harlow, S. M. Till, T. A. Russell, L. S. Wijetunge, P. Kind, and A. Contractor, “Critical period plasticity is disrupted in the barrel cortex of FMR1 knockout mice,” Neuron 65, no. 3 (2010): 385–398. 10.1016/j.neuron.2010.01.024.

10. G. Curia, T. Papouin, P. Séguéla, and M. Avoli, “Downregulation of tonic GABAergic inhibition in a mouse model of fragile X syndrome,” Cerebral Cortex 19, no. 7 (2009): 1515–1520. 10.1093/cercor/bhn159.

11. P. W. Frankland, Y. Wang, B. Rosner, et al., “Sensorimotor gating abnormalities in young males with fragile X syndrome and Fmr1-knockout mice,” Molecular Psychiatry 9, no. 4 (2004): 417–425. 10.1038/sj.mp.4001432.

12. S. L. Wadle, T. C. Ritter, T. T. X. Wadle, and J. J. Hirtz, “Topography and ensemble activity in the auditory cortex of a mouse model of fragile X syndrome,” eNeuro 11, no. 5 (2024): ENEURO.0396-23.2024. 10.1523/eneuro.0396-23.2024.

13. S. S. Kornfeld-Sylla, C. Gelegen, J. E. Norris, et al., “A human electrophysiological signature of Fragile X pathophysiology is shared in V1 of Fmr1-/y mice,” Nature Communications 17, no. 1 (2026): 1497. 10.1038/s41467-026-69243-0.

14. G. M. Shepherd, “Corticostriatal connectivity and its role in disease,” Nature Reviews Neuroscience 14, no. 4 (2013): 278–291. 10.1038/nrn3469.

15. H. Hintiryan, N. N. Foster, I. Bowman, et al., “The mouse cortico-striatal projectome,” Nature Neuroscience 19, no. 8 (2016): 1100–1114. 10.1038/nn.4332.

16. B. J. Hunnicutt, B. C. Jongbloets, W. T. Birdsong, K. J. Gertz, H. Zhong, and T. Mao, “A comprehensive excitatory input map of the striatum reveals novel functional organization,” eLife 5 (2016): e19103. 10.7554/elife.19103.

17. L. A. Gunaydin, L. Grosenick, J. C. Finkelstein, et al., “Natural neural projection dynamics underlying social behavior,” Cell 157, no. 7 (2014): 1535–1551. 10.1016/j.cell.2014.05.017.

18. J. Peça, C. Feliciano, J. T. Ting, et al., “Shank3 mutant mice display autistic-like behaviours and striatal dysfunction,” Nature 472, no. 7344 (2011): 437–442. 10.1038/nature09965.

19. E. Burguière, P. Monteiro, G. Feng, and A. M. Graybiel, “Optogenetic stimulation of lateral orbitofronto-striatal pathway suppresses compulsive behaviors,” Science 340, no. 6137 (2013): 1243–1246. 10.1126/science.1232380.

20. N. Barnea-Goraly, S. Eliez, M. Hedeus, et al., “White matter tract alterations in fragile X syndrome: preliminary evidence from diffusion tensor imaging,” American Journal of Medical Genetics Part B: Neuropsychiatric Genetics 118B, no. 1 (2003): 81–88. 10.1002/ajmg.b.10035.

21. B. W. Haas, N. Barnea-Goraly, A. A. Lightbody, et al., “Early white-matter abnormalities of the ventral frontostriatal pathway in fragile X syndrome,” Developmental Medicine & Child Neurology 51, no. 8 (2009): 593–599. 10.1111/j.1469-8749.2009.03295.x.

22. J. L. Bruno, S. M. H. Hosseini, M. Saggar, E. M. Quintin, M. M. Raman, and A. L. Reiss, “Altered brain network segregation in fragile X syndrome revealed by structural connectomics,” Cerebral Cortex 27, no. 3 (2017): 2249–2259. 10.1093/cercor/bhw055.

23. M. R. Swanson, J. J. Wolff, M. D. Shen, et al., “Development of white matter circuitry in infants with fragile X syndrome,” JAMA Psychiatry 75, no. 5 (2018): 505–513. 10.1001/jamapsychiatry.2018.0180.

24. S. S. Hall, R. F. Dougherty, and A. L. Reiss, “Profiles of aberrant white matter microstructure in fragile X syndrome,” NeuroImage: Clinical 11 (2016): 133–138. 10.1016/j.nicl.2016.01.013.

25. K. Yamamuro, M. Ikehara, Y. Noriyama, et al., “Oxytocin receptor neurons in the paraventricular thalamus as a nexus for social behaviour and fear,” Brain 149, no. 6 (2026): 2151–2165. 10.1093/brain/awaf421.

26. C. Qi, W. Sima, H. Mao, et al., “Anterior cingulate cortex parvalbumin and somatostatin interneurons shape social behavior in male mice,” Nature Communications 16, no. 1 (2025): 4156. 10.1038/s41467-025-59473-z.

27. T. T. Hsu, T. N. Huang, C. Y. Wang, and Y. P. Hsueh, “Deep brain stimulation of the Tbr1-deficient mouse model of autism spectrum disorder at the basolateral amygdala alters amygdalar connectivity, whole-brain synchronization, and social behaviors,” PLOS Biology 22, no. 7 (2024): e3002646. 10.1371/journal.pbio.3002646.

28. M. Prieto, A. Folci, G. Poupon, et al., “Missense mutation of Fmr1 results in impaired AMPAR-mediated plasticity and socio-cognitive deficits in mice,” Nature Communications 12, no. 1 (2021): 1557. 10.1038/s41467-021-21820-1.

29. L. K. Myrick, P. Y. Deng, H. Hashimoto, et al., “Independent role for presynaptic FMRP revealed by an FMR1 missense mutation associated with intellectual disability and seizures,” Proceedings of the National Academy of Sciences 112, no. 4 (2015): 949–956. 10.1073/pnas.1423094112.

30. Z. Okray, C. E. de Esch, H. Van Esch, et al., “A novel fragile X syndrome mutation reveals a conserved role for the carboxy-terminus in FMRP localization and function,” EMBO Molecular Medicine 7, no. 4 (2015): 423–437. 10.15252/emmm.201404576.

31. L. K. Myrick, M. Nakamoto-Kinoshita, N. M. Lindor, S. Kirmani, X. Cheng, and S. T. Warren, “Fragile X syndrome due to a missense mutation,” European Journal of Human Genetics 22, no. 10 (2014): 1185–1189. 10.1038/ejhg.2013.311.

32. M. Harbers, Z. Wei, H. Nakao, et al., “FMR1 mutant marmosets show fragile X syndrome phenotypes,” Cell Reports 44, no. 9 (2025): 116208. 10.1016/j.celrep.2025.116208.

33. P. Licznerski, H. A. Park, H. Rolyan, et al., “ATP synthase c-subunit leak causes aberrant cellular metabolism in fragile X syndrome,” Cell 182, no. 5 (2020): 1170–1185.e9. 10.1016/j.cell.2020.07.008.

34. M. Shen, C. L. Sirois, Y. Guo, et al., “Species-specific FMRP regulation of RACK1 is critical for prenatal cortical development,” Neuron 111, no. 24 (2023): 3988–4005.e11. 10.1016/j.neuron.2023.09.014.

35. M. Shen, F. Wang, M. Li, et al., “Reduced mitochondrial fusion and Huntingtin levels contribute to impaired dendritic maturation and behavioral deficits in Fmr1-mutant mice,” Nature Neuroscience 22, no. 3 (2019): 386–400. 10.1038/s41593-019-0338-y.

36. A. R. Fenton, R. Peng, C. Bond, et al., “FMRP regulates MFF translation to locally direct mitochondrial fission in neurons,” Nature Cell Biology 26, no. 12 (2024): 2061–2074. 10.1038/s41556-024-01544-2.

37. R. Z. Yang, Y. Wang, Z. Wu, et al., “Ambient light alleviates retinal neurodegeneration in mice by powering mitochondria via the engineered optoenergetic rhodopsin,” Signal Transduction and Targeted Therapy 10, no. 1 (2025): 355. 10.1038/s41392-025-02450-1.

38. M. D. C. Marín, M. Konno, H. Yawo, and K. Inoue, “Converting a natural-light-driven outward proton pump rhodopsin into an artificial inward proton pump,” Journal of the American Chemical Society 145, no. 20 (2023): 10938–10942. 10.1021/jacs.2c12602.

39. X. Li, P. S. Morgan, J. Ashburner, J. Smith, and C. Rorden, “The first step for neuroimaging data analysis: DICOM to NIfTI conversion,” Journal of Neuroscience Methods 264 (2016): 47–56. 10.1016/j.jneumeth.2016.03.001.

40. J. D. Tournier, R. Smith, D. Raffelt, et al., “MRtrix3: a fast, flexible and open software framework for medical image processing and visualisation,” NeuroImage 202 (2019): 116137. 10.1016/j.neuroimage.2019.116137.

41. M. Jenkinson, C. F. Beckmann, T. E. Behrens, M. W. Woolrich, and S. M. Smith, “FSL,” NeuroImage 62, no. 2 (2012): 782–790. 10.1016/j.neuroimage.2011.09.015.

42. J. Veraart, D. S. Novikov, D. Christiaens, B. Ades-Aron, J. Sijbers, and E. Fieremans, “Denoising of diffusion MRI using random matrix theory,” NeuroImage 142 (2016): 394–406. 10.1016/j.neuroimage.2016.08.016.

43. E. Kellner, B. Dhital, V. G. Kiselev, and M. Reisert, “Gibbs-ringing artifact removal based on local subvoxel-shifts,” Magnetic Resonance in Medicine 76, no. 5 (2016): 1574–1581. 10.1002/mrm.26054.

44. J. L. R. Andersson and S. N. Sotiropoulos, “An integrated approach to correction for off-resonance effects and subject movement in diffusion MR imaging,” NeuroImage 125 (2016): 1063–1078. 10.1016/j.neuroimage.2015.10.019.

45. N. J. Tustison, B. B. Avants, P. A. Cook, et al., “N4ITK: improved N3 bias correction,” IEEE Transactions on Medical Imaging 29, no. 6 (2010): 1310–1320. 10.1109/tmi.2010.2046908.

46. S. M. Smith, M. Jenkinson, H. Johansen-Berg, et al., “Tract-based spatial statistics: voxelwise analysis of multi-subject diffusion data,” NeuroImage 31, no. 4 (2006): 1487–1505. 10.1016/j.neuroimage.2006.02.024.

47. S. Mori, K. Oishi, H. Jiang, et al., “Stereotaxic white matter atlas based on diffusion tensor imaging in an ICBM template,” NeuroImage 40, no. 2 (2008): 570–582. 10.1016/j.neuroimage.2007.12.035.

48. R. E. Smith, J. D. Tournier, F. Calamante, and A. Connelly, “Anatomically-constrained tractography: improved diffusion MRI streamlines tractography through effective use of anatomical information,” NeuroImage 62, no. 3 (2012): 1924–1938. 10.1016/j.neuroimage.2012.06.005.

49. R. E. Smith, J. D. Tournier, F. Calamante, and A. Connelly, “SIFT2: enabling dense quantitative assessment of brain white matter connectivity using streamlines tractography,” NeuroImage 119 (2015): 338–351. 10.1016/j.neuroimage.2015.06.092.

50. B. Fischl, D. H. Salat, E. Busa, et al., “Whole brain segmentation: automated labeling of neuroanatomical structures in the human brain,” Neuron 33, no. 3 (2002): 341–355. 10.1016/S0896-6273(02)00569-X.

51. R. S. Desikan, F. Ségonne, B. Fischl, et al., “An automated labeling system for subdividing the human cerebral cortex on MRI scans into gyral based regions of interest,” NeuroImage 31, no. 3 (2006): 968–980. 10.1016/j.neuroimage.2006.01.021.

52. Q. Wang, S. L. Ding, Y. Li, et al., “The Allen Mouse Brain Common Coordinate Framework: a 3D reference atlas,” Cell 181, no. 4 (2020): 936–953.e20. 10.1016/j.cell.2020.04.007.

53. N. C. Klapoetke, Y. Murata, S. S. Kim, et al., “Independent optical excitation of distinct neural populations,” Nature Methods 11, no. 3 (2014): 338–346. 10.1038/nmeth.2836.

54. V. Gradinaru, F. Zhang, C. Ramakrishnan, et al., “Molecular and cellular approaches for diversifying and extending optogenetics,” Cell 141, no. 1 (2010): 154–165. 10.1016/j.cell.2010.02.037.

55. X. Zhang, J. Zhang, and M. Shen, “MitoQ attenuates maternal immune activation-induced mitochondrial and ER stress programs and ameliorates psychiatric-like behavioral deficits,” Free Radical Biology and Medicine 253 (2026): 351–363. 10.1016/j.freeradbiomed.2026.05.326.

56. M. Shao, Y. Wu, H. Wang, et al., “Counterbalancing O-GlcNAcylation and STAT3 phosphorylation in ventral tegmental area dopaminergic neurons mediates behavioral adaptations to acute restraint stress,” Advanced Science 12, no. 39 (2025): e02701. 10.1002/advs.202502701.

57. M. Shen, Y. Guo, Q. Dong, et al., “FXR1 regulation of parvalbumin interneurons in the prefrontal cortex is critical for schizophrenia-like behaviors,” Molecular Psychiatry 26, no. 11 (2021): 6845–6867. 10.1038/s41380-021-01096-z.

58. G. Cui, S. B. Jun, X. Jin, et al., “Deep brain optical measurements of cell type-specific neural activity in behaving mice,” Nature Protocols 9, no. 6 (2014): 1213–1228. 10.1038/nprot.2014.080.

59. T. W. Chen, T. J. Wardill, Y. Sun, et al., “Ultrasensitive fluorescent proteins for imaging neuronal activity,” Nature 499, no. 7458 (2013): 295–300. 10.1038/nature12354.

60. J. Zhou, H. Wang, R. Shen, et al., “Mitochondrial-targeted antioxidant MitoQ provides neuroprotection and reduces neuronal apoptosis in experimental traumatic brain injury possibly via the Nrf2-ARE pathway,” American Journal of Translational Research 10, no. 6 (2018): 1887–1899.

61. D. Kim, J. M. Paggi, C. Park, C. Bennett, and S. L. Salzberg, “Graph-based genome alignment and genotyping with HISAT2 and HISAT-genotype,” Nature Biotechnology 37, no. 8 (2019): 907–915. 10.1038/s41587-019-0201-4.

62. Y. Liao, G. K. Smyth, and W. Shi, “featureCounts: an efficient general purpose program for assigning sequence reads to genomic features,” Bioinformatics 30, no. 7 (2014): 923–930. 10.1093/bioinformatics/btt656.

63. M. I. Love, W. Huber, and S. Anders, “Moderated estimation of fold change and dispersion for RNA-seq data with DESeq2,” Genome Biology 15, no. 12 (2014): 550. 10.1186/s13059-014-0550-8.

64. A. Subramanian, P. Tamayo, V. K. Mootha, et al., “Gene set enrichment analysis: a knowledge-based approach for interpreting genome-wide expression profiles,” Proceedings of the National Academy of Sciences 102, no. 43 (2005): 15545–15550. 10.1073/pnas.0506580102.

65. A. Liberzon, C. Birger, H. Thorvaldsdóttir, M. Ghandi, J. P. Mesirov, and P. Tamayo, “The Molecular Signatures Database hallmark gene set collection,” Cell Systems 1, no. 6 (2015): 417–425. 10.1016/j.cels.2015.12.004.

66. G. Yu, L. G. Wang, Y. Han, and Q. Y. He, “clusterProfiler: an R package for comparing biological themes among gene clusters,” OMICS: A Journal of Integrative Biology 16, no. 5 (2012): 284–287. 10.1089/omi.2011.0118.

67. M. J. Hurley, R. M. J. Deacon, A. W. E. Chan, D. Baker, D. L. Selwood, and P. Cogram, “Reversal of behavioural phenotype by the cannabinoid-like compound VSN16R in fragile X syndrome mice,” Brain 145, no. 1 (2022): 76–82. 10.1093/brain/awab246.

68. K. Chalkiadaki, M. Hooshmandi, G. Lach, et al., “Mnk1/2 kinases regulate memory and autism-related behaviours via Syngap1,” Brain 146, no. 5 (2023): 2175–2190. 10.1093/brain/awac398.

69. K. Zhang, Y. J. Li, Y. Guo, et al., “Elevated progranulin contributes to synaptic and learning deficit due to loss of fragile X mental retardation protein,” Brain 140, no. 12 (2017): 3215–3232. 10.1093/brain/awx265.

70. M. Pagani, V. Zerbi, S. Gini, et al., “Autism subtypes identified using cross-species functional connectivity analyses,” Nature Neuroscience 29, no. 6 (2026): 1476–1487. 10.1038/s41593-026-02287-z.

71. R. T. Peixoto, W. Wang, D. M. Croney, Y. Kozorovitskiy, and B. L. Sabatini, “Early hyperactivity and precocious maturation of corticostriatal circuits in Shank3B(-/-) mice,” Nature Neuroscience 19, no. 5 (2016): 716–724. 10.1038/nn.4260.

72. F. Longo, S. Aryal, P. G. Anastasiades, et al., “Cell-type-specific disruption of cortico-striatal circuitry drives repetitive patterns of behavior in fragile X syndrome model mice,” Cell Reports 42, no. 8 (2023): 112901. 10.1016/j.celrep.2023.112901.

73. G. Giua, O. Lassalle, L. Makrini-Maleville, E. Valjent, P. Chavis, and O. J. J. Manzoni, “Investigating cell-specific effects of FMRP deficiency on spiny projection neurons in a mouse model of Fragile X syndrome,” Frontiers in Cellular Neuroscience 17 (2023): 1146647. 10.3389/fncel.2023.1146647.

74. J. L. Huebschman, C. A. Monterrey, D. M. Foster, et al., “The role of the dorsal striatum in a mouse model for fragile X syndrome: behavioral and dendritic spine assessment,” Brain Research 1795 (2022): 148060. 10.1016/j.brainres.2022.148060.

75. A. A. Legaria, B. A. Matikainen-Ankney, B. Yang, et al., “Fiber photometry in striatum reflects primarily nonsomatic changes in calcium,” Nature Neuroscience 25, no. 9 (2022): 1124–1128. 10.1038/s41593-022-01152-z.

