## Supplementary Material for "FMR1 Perturbations Disrupt Sensory Corticostriatal Circuits and Mitochondrial Energetics in Fragile X Syndrome"

### These authors contributed equally.

#### Supporting Information

Supplementary Table 1. Key resources

Supplementary Table 2. Statistical analyses

Supplementary Table 3. Summary of whole-brain white matter microstructural alterations in FXS compared to TD

Supplementary Figure 1. Age and sex distributions in the human cohorts.

Supplementary Figure 2. Increased radial diffusivity in participants with fragile X syndrome.

Supplementary Figure 3. Fractional anisotropy in additional S1–caudoputamen and S1–striatum pathways.

Supplementary Figure 4. S1 calcium responses to local optogenetic stimulation in a nonsocial condition.

Supplementary Figure 5. DMS calcium responses to S1 optogenetic stimulation in *Fmr1*-deficient mice.

Supplementary Figure 6. *Fmr1*-I234T expression and S1–DMS projection activation modify behavioural phenotypes.

Supplementary Figure 7. Transcriptomic structure and OXPHOS pathway scores from regional RNA-sequencing analysis.

#### Supplementary Table 1. Key resources

| REAGENT or RESOURCE | SOURCE | IDENTIFIER |
|---|---|---|
| **Antibodies** | | |
| mouse anti-NeuN | Sigma-Aldrich | MAB377 |
| rabbit anti-PGC-1α/β | Abcam | ab54481 |
| goat anti-mouse Alexa Fluor 488 | Thermo Fisher Scientific | A11029 |
| goat anti-rabbit Alexa Fluor 647 | Thermo Fisher Scientific | A21245 |
| **Critical commercial assays** | | |
| Rhodamine 800 | Sigma-Aldrich | CAS# 137993-41-0 |
| Transferrin | Sigma-Aldrich | CAS# 11096-37-0 |
| Cytarabine | Sigma-Aldrich | CAS# 147-94-4 |
| B27 supplement | Invitrogen | Cat# 17504044 |
| Matrigel | BD Biosciences | Cat# 354248 |
| FastPure Complex Tissue/Cell Total RNA Isolation Kit | Vazyme | RC113-01 |
| MitoQ | Selleck Chemicals | S8978 |
| **Virus** | | |
| rAAV-hSyn-axon-GCaMP6m-WPRE-hGH polyA | BrainVTA | Cat# PT-1224; AAV2/9 |
| rAAV-hSyn-ChrimsonR-tdTomato-WPRE-bGH polyA | BrainVTA | Cat# PT-1432; AAV2/9 |
| rAAV-hSyn-GCaMP6m-WPRE-hGH polyA | BrainVTA | Cat# PT-0148; AAV2/9 |
| rAAV-CaMKIIα-GCaMP6m-WPRE-hGH polyA | BrainVTA | Cat# PT-0111; AAV2/9 |
| rAAV-CaMKIIα-ChrimsonR-mCherry-WPREs | BrainVTA | Cat# PT-4285; AAV2/9 |
| rAAV-EF1α-DIO-GCaMP6m-WPRE-hGH polyA | BrainVTA | Cat# PT-0283; AAV2/9 |
| rAAV-hSyn-CRE-WPRE-hGH polyA | BrainVTA | Cat# PT-0136; AAV2/R |
| rAAV-hSyn-DIO-ChrimsonR-mCherry-WPRE-hGH polyA | BrainVTA | Cat# PT-1374; AAV2/9 |
| rAAV-EF1α-DIO-eNpHR3.0-EYFP-WPRE-hGH polyA | BrainVTA | Cat# PT-0006; AAV2/9 |
| rAAV-hSyn-mt-EcGAPR-EGFP | BrainCase | AAV9 |
| pAAV-hSyn-*Fmr1*(WT)-3×FLAG-P2A-GFP | Vigene Biosciences | AAV9 |

| REAGENT or RESOURCE | SOURCE | IDENTIFIER |
| --- | --- | --- |
| pAAV-hSyn-*Fmr1*(I234T)-3×FLAG-P2A-GFP | Vigene Biosciences | AAV9 |
| pAAV-hSyn-*Fmr1*(E329G)-3×FLAG-P2A-GFP | Vigene Biosciences | AAV9 |
| pAAV-hSyn-*Fmr1*(R586Q)-3×FLAG-P2A-GFP | Vigene Biosciences | AAV9 |
| pAAV-hSyn-GFP | Vigene Biosciences | AAV9 |
| **Animals** | | |
| Mouse: *Fmr1* knockout | The Jackson Laboratory | Stock no. 004624 |
| **Deposited Data** | | |
| RNA-seq data | | GEO: GSE342691 |
| **Software and algorithms** | | |
| MRtrix3 | MRtrix3 | v3.0.8 |
| FSL | FMRIB Software Library | v6.0.7.22 |
| MATLAB | MathWorks | R2017b |
| ImageJ | NIH | 1.54g |
| GraphPad Prism | GraphPad Software | 9.5.1 |
| SPSS | IBM | 22 |

#### Supplementary Table 2. Statistical analyses

| Figure / Panel and endpoint | Sample size (*n*) | Normality test | Variance test | Analysis | *P* Value | Degrees of freedom and test statistic |
|---|---|---|---|---|---|---|
| **Figure 1** | | | | | | |
| Figure 1D<br>L postcentral–L caudate SIFT2 | FXS = 52;<br>TD = 64 | Shapiro-Wilk test, P(FXS) < 0.0001; P(TD) < 0.0001 | Bartlett's test: P < 0.0001 | unpaired two-tailed t-test | P = 0.0082 (**) | t(114) = 2.6897 |
| Figure 1D<br>L precentral–L caudate SIFT2 | FXS = 52;<br>TD = 64 | Shapiro-Wilk test, P(FXS) = 0.4265; P(TD) = 0.1811 | Bartlett's test: P = 0.0022 | unpaired two-tailed t-test | P = 0.0031 (**) | t(114) = 3.0249 |
| Figure 1D<br>L paracentral–L pallidum SIFT2 | FXS = 52;<br>TD = 64 | Shapiro-Wilk test, P(FXS) < 0.0001; P(TD) < 0.0001 | Bartlett's test: P < 0.0001 | unpaired two-tailed t-test | P = 0.0382 (*) | t(114) = -2.0966 |
| Figure 1D<br>L paracentral–L putamen SIFT2 | FXS = 52;<br>TD = 64 | Shapiro-Wilk test, P(FXS) < 0.0001; P(TD) < 0.0001 | Bartlett's test: P = 0.0117 | unpaired two-tailed t-test | P = 0.0233 (*) | t(114) = -2.2998 |
| Figure 1D<br>L supramarginal–R caudate SIFT2 | FXS = 52;<br>TD = 64 | Shapiro-Wilk test, P(FXS) < 0.0001; P(TD) < 0.0001 | Bartlett's test: P < 0.0001 | unpaired two-tailed t-test | P = 0.0469 (*) | t(114) = -2.009 |
| **Figure 2** | | | | | | |
| Figure 2D<br>Mouth FA | WT = 7;<br>Fmr1-KO = 7 | Shapiro-Wilk test, P(WT) = 0.4143; P(Fmr1-KO) = 0.0585 | Bartlett's test: P = 0.7718 | unpaired two-tailed t-test | P = 0.0412 (*) | t(12) = 2.2867 |
| Figure 2D<br>Upper limb FA | WT = 7;<br>Fmr1-KO = 7 | Shapiro-Wilk test, P(WT) = 0.4424; P(Fmr1-KO) = 0.3405 | Bartlett's test: P = 0.4593 | unpaired two-tailed t-test | P = 0.0269 (*) | t(12) = 2.5192 |
| Figure 2F<br>Mouth FA | WT = 7;<br>Fmr1-KO = 7 | Shapiro-Wilk test, P(WT) = 0.5231; P(Fmr1-KO) = 0.8225 | Bartlett's test: P = 0.5495 | unpaired two-tailed t-test | P = 0.0171 (*) | t(12) = 2.7646 |
| Figure 2F<br>Upper limb FA | WT = 7;<br>Fmr1-KO = 7 | Shapiro-Wilk test, P(WT) = 0.7703; P(Fmr1-KO) = 0.9094 | Bartlett's test: P = 0.2527 | unpaired two-tailed t-test | P = 0.1937 (ns) | t(12) = 1.3770 |
| **Figure 3** | | | | | | |
| Figure 3C<br>Novel location recognition index | WT = 16;<br>Fmr1-KO = 14 | Shapiro-Wilk test, P(WT) = 0.0725; P(Fmr1-KO) = 0.9254 | Bartlett's test: P = 0.2009 | unpaired two-tailed t-test | P < 0.0001 (****) | t(28) = 4.6049 |

| Figure / Panel and endpoint | Sample size (*n*) | Normality test | Variance test | Analysis | *P* Value | Degrees of freedom and test statistic |
|---|---|---|---|---|---|---|
| Figure 3D Object-contact AUC | WT = 9; Fmr1-KO = 9 | Shapiro-Wilk test, P(WT) = 0.3121; P(Fmr1-KO) = 0.4451 | Bartlett's test: P = 0.2349 | unpaired two-tailed t-test | P = 0.2963 (ns) | t(16) = -1.0797 |
| Figure 3E Object-contact AUC | WT = 6; Fmr1-KO = 6 | Shapiro-Wilk test, P(WT) = 0.5238; P(Fmr1-KO) = 0.4653 | Bartlett's test: P = 0.3336 | unpaired two-tailed t-test | P = 0.0472 (*) | t(10) = -2.2620 |
| Figure 3F Object-contact AUC | WT = 8; Fmr1-KO = 8 | Shapiro-Wilk test, P(WT) = 0.4799; P(Fmr1-KO) = 0.3884 | Bartlett's test: P = 0.9986 | unpaired two-tailed t-test | P = 0.0956 (ns) | t(14) = -1.7872 |
| Figure 3G Object-contact AUC | WT = 8; Fmr1-KO = 7 | Shapiro-Wilk test, P(WT) = 0.9834; P(Fmr1-KO) = 0.6399 | Bartlett's test: P = 0.2918 | unpaired two-tailed t-test | P = 0.0011 (**) | t(13) = -4.1690 |
| Figure 3J Sociability preference index | WT = 16; Fmr1-KO = 15 | Shapiro-Wilk test, P(WT) = 0.1080; P(Fmr1-KO) = 0.5198 | Bartlett's test: P = 0.2308 | unpaired two-tailed t-test | P = 0.0099 (**) | t(29) = 2.7597 |
| Figure 3M Social novelty preference index | WT = 16; Fmr1-KO = 15 | Shapiro-Wilk test, P(WT) = 0.4417; P(Fmr1-KO) = 0.4583 | Bartlett's test: P = 0.5404 | unpaired two-tailed t-test | P = 0.0269 (*) | t(29) = 2.3319 |
| Figure 3N Stranger-contact AUC | WT = 9; Fmr1-KO = 9 | Shapiro-Wilk test, P(WT) = 0.7901; P(Fmr1-KO) = 0.0194 | Bartlett's test: P = 0.7605 | Mann–Whitney U test | P = 0.0056 (**) | U = 10.0000 |
| Figure 3O Stranger-contact AUC | WT = 6; Fmr1-KO = 6 | Shapiro-Wilk test, P(WT) = 0.0344; P(Fmr1-KO) = 0.7417 | Bartlett's test: P = 0.0617 | Mann–Whitney U test | P = 0.0022 (**) | U = 0.0000 |
| Figure 3P Stranger-contact AUC | WT = 8; Fmr1-KO = 8 | Shapiro-Wilk test, P(WT) = 0.4290; P(Fmr1-KO) = 0.0562 | Bartlett's test: P = 0.5398 | unpaired two-tailed t-test | P = 0.0312 (*) | t(14) = 2.3943 |
| Figure 3Q Stranger-contact AUC | WT = 8; Fmr1-KO = 7 | Shapiro-Wilk test, P(WT) = 0.8333; P(Fmr1-KO) = 0.8384 | Bartlett's test: P = 0.0268 | Welch's unpaired two-tailed t-test | P = 0.0293 (*) | t(9.1204) = 2.5811 |
| **Figure 4** | | | | | | |
| Figure 4D Fmr1-KO sociability preference index | n = 7 paired observations | Shapiro-Wilk test of paired differences, P = 0.4782 | Not applicable (paired design) | paired two-tailed t-test | P = 0.0310 (*) | t(6) = -2.8038 |

| Figure / Panel and endpoint | Sample size (*n*) | Normality test | Variance test | Analysis | *P* Value | Degrees of freedom and test statistic |
|---|---|---|---|---|---|---|
| Figure 4H<br>WT sociability preference index | n = 9 paired observations | Shapiro-Wilk test of paired differences, P = 0.8477 | Not applicable (paired design) | paired two-tailed t-test | P = 0.0039 (**) | t(8) = 4.0131 |
| Figure 4K<br>Novel location recognition index | GFP = 9;<br>I234T = 9;<br>E329G = 6;<br>R586Q = 6 | Shapiro-Wilk test, P(GFP) = 0.4123; P(I234T) = 0.5909; P(E329G) = 0.2647; P(R586Q) = 0.1423 | Bartlett's test: P = 0.0582 | one-way ANOVA with Dunnett's multiple-comparisons test | Omnibus P = 0.0061 (**)<br>Post hoc:<br>GFP vs I234T: P = 0.0274 (*)<br>GFP vs E329G: P = 0.5352 (ns)<br>GFP vs R586Q: P = 0.4334 (ns) | F(3, 26) = 5.1920 |
| Figure 4M<br>Sociability preference index | GFP = 9;<br>I234T = 8;<br>E329G = 7;<br>R586Q = 6 | Shapiro-Wilk test, P(GFP) = 0.1263; P(I234T) = 0.7271; P(E329G) = 0.2729; P(R586Q) = 0.6669 | Bartlett's test: P = 0.1599 | one-way ANOVA with Dunnett's multiple-comparisons test | Omnibus P = 0.0014 (**)<br>Post hoc:<br>GFP vs I234T: P = 0.0204 (*)<br>GFP vs E329G: P = 0.000999 (***)<br>GFP vs R586Q: P = 0.0062 (**) | F(3, 26) = 6.9165 |
| Figure 4P<br>I234T sociability preference index | n = 12 paired observations | Shapiro-Wilk test of paired differences, P = 0.9887 | Not applicable (paired design) | paired two-tailed t-test | P = 0.0014 (**) | t(11) = -4.2317 |
| Figure 4P<br>E329G sociability preference index | n = 10 paired observations | Shapiro-Wilk test of paired differences, P = 0.5209 | Not applicable (paired design) | paired two-tailed t-test | P = 0.0018 (**) | t(9) = -4.3568 |
| Figure 4P<br>R586Q sociability preference index | n = 8 paired observations | Shapiro-Wilk test of paired differences, P = 0.2516 | Not applicable (paired design) | paired two-tailed t-test | P = 0.0143 (*) | t(7) = -3.2374 |

| **Figure / Panel and endpoint** | **Sample size (*n*)** | **Normality test** | **Variance test** | **Analysis** | ***P* Value** | **Degrees of freedom and test statistic** |
|---|---|---|---|---|---|---|
| **Figure 6** | | | | | | |
| Figure 6B Novel-location discrimination index | Vehicle WT = 7; Vehicle Fmr1-KO = 7; MitoQ WT = 9; MitoQ Fmr1-KO = 12 | Shapiro-Wilk test, P(Vehicle WT) = 0.3891; P(Vehicle Fmr1-KO) = 0.3530; P(MitoQ WT) = 0.1618; P(MitoQ Fmr1-KO) = 0.1803 | Bartlett's test: P = 0.0612 | two-way ANOVA with Tukey's multiple-comparisons test | Genotype: P < 0.0001 (****) Treatment: P = 0.3324 (ns) Genotype × treatment: P = 0.0024 (**) Tukey: Vehicle WT vs Vehicle Fmr1-KO: P < 0.0001 (****) Vehicle WT vs MitoQ WT: P = 0.3934 (ns) Vehicle WT vs MitoQ Fmr1-KO: P = 0.0013 (**) Vehicle Fmr1-KO vs MitoQ WT: P < 0.0001 (****) Vehicle Fmr1-KO vs MitoQ Fmr1-KO: P = 0.0188 (*) MitoQ WT vs MitoQ Fmr1-KO: P = 0.0581 (ns) | Genotype: F(1, 31) = 44.6238 Treatment: F(1, 31) = 0.9698 Genotype × treatment: F(1, 31) = 10.9702 |

| Figure / Panel and endpoint | Sample size (*n*) | Normality test | Variance test | Analysis | *P* Value | Degrees of freedom and test statistic |
|---|---|---|---|---|---|---|
| Figure 6C Social-interaction discrimination index | Vehicle WT = 7; Vehicle Fmr1-KO = 7; MitoQ WT = 9; MitoQ Fmr1-KO = 12 | Shapiro-Wilk test, P(Vehicle WT) = 0.0485; P(Vehicle Fmr1-KO) = 0.8362; P(MitoQ WT) = 0.2007; P(MitoQ Fmr1-KO) = 0.6218 | Bartlett's test: P = 0.6977 | two-way ANOVA with Tukey's multiple-comparisons test | Genotype: P = 0.0016 (**) Treatment: P = 0.0006 (***) Genotype × treatment: P = 0.1074 (ns) Tukey: Vehicle WT vs Vehicle Fmr1-KO: P = 0.0120 (*) Vehicle WT vs MitoQ WT: P = 0.4669 (ns) Vehicle WT vs MitoQ Fmr1-KO: P = 0.9943 (ns) Vehicle Fmr1-KO vs MitoQ WT: P = 0.0001 (***) Vehicle Fmr1-KO vs MitoQ Fmr1-KO: P = 0.0021 (**) MitoQ WT vs MitoQ Fmr1-KO: P = 0.5028 (ns) | Genotype: F(1, 31) = 11.9204 Treatment: F(1, 31) = 14.4160 Genotype × treatment: F(1, 31) = 2.7493 |

| Figure / Panel and endpoint | Sample size (*n*) | Normality test | Variance test | Analysis | *P* Value | Degrees of freedom and test statistic |
|---|---|---|---|---|---|---|
| Figure 6D Social-novelty discrimination index | Vehicle WT = 7; Vehicle Fmr1-KO = 7; MitoQ WT = 9; MitoQ Fmr1-KO = 12 | Shapiro-Wilk test, P(Vehicle WT) = 0.6897; P(Vehicle Fmr1-KO) = 0.4067; P(MitoQ WT) = 0.9814; P(MitoQ Fmr1-KO) = 0.6520 | Bartlett's test: P = 0.0819 | two-way ANOVA with Tukey's multiple-comparisons test | Genotype: P < 0.0001 (****) Treatment: P < 0.0001 (****) Genotype × treatment: P < 0.0001 (****) Tukey: Vehicle WT vs Vehicle Fmr1-KO: P < 0.0001 (****) Vehicle WT vs MitoQ WT: P = 0.9655 (ns) Vehicle WT vs MitoQ Fmr1-KO: P = 0.9789 (ns) Vehicle Fmr1-KO vs MitoQ WT: P < 0.0001 (****) Vehicle Fmr1-KO vs MitoQ Fmr1-KO: P < 0.0001 (****) MitoQ WT vs MitoQ Fmr1-KO: P = 0.9995 (ns) | Genotype: F(1, 31) = 28.6332 Treatment: F(1, 31) = 23.1382 Genotype × treatment: F(1, 31) = 30.1533 |
| Figure 6F Transformed Rhodamine 800 endpoint change | EGFP control = 6; mt-EcGAPR = 9 | Shapiro-Wilk test, P(EGFP control) = 0.1992; P(mt-EcGAPR) = 0.4961 | Bartlett's test: P = 0.5566 | unpaired two-tailed t-test | P = 0.0040 (**) | t(13) = -3.4854 |
| Figure 6H Rhodamine 800 area under curve | EGFP control = 9; mt-EcGAPR = 10 | Shapiro-Wilk test, P(EGFP control) = 0.3004; P(mt-EcGAPR) = 0.1716 | Bartlett's test: P = 0.9669 | unpaired two-tailed t-test | P = 0.0380 (*) | t(17) = -2.2503 |
| Figure 6K S1 PGC-1α immunofluorescence | Mito-OptoEA− = 9 sections; Mito-OptoEA+ = 8 sections; 3 mice per group | Shapiro-Wilk test, P(Mito-OptoEA−) = 0.2619; P(Mito-OptoEA+) = 0.3750 | Bartlett's test: P = 0.5375 | unpaired two-tailed t-test | P = 0.0081 (**) | t(15) = -3.0529 |

| Figure / Panel and endpoint | Sample size (*n*) | Normality test | Variance test | Analysis | *P* Value | Degrees of freedom and test statistic |
|---|---|---|---|---|---|---|
| Figure 6K<br>DMS PGC-1α immunofluorescence | Mito-OptoEA− = 9 sections;<br>Mito-OptoEA+ = 8 sections;<br>3 mice per group | Shapiro-Wilk test, P(Mito-OptoEA−) = 0.3726; P(Mito-OptoEA+) = 0.5241 | Bartlett's test: P = 0.5405 | unpaired two-tailed t-test | P = 0.0041 (**) | t(15) = -3.3831 |
| Figure 6L<br>Fmr1-KO novel location recognition index | n = 10 paired observations | Shapiro-Wilk test of paired differences, P = 0.7241 | Not applicable (paired design) | paired two-tailed t-test | P = 0.0443 (*) | t(9) = -2.3365 |
| Figure 6M<br>WT+Fmr1-I234T novel location recognition index | n = 10 paired observations | Shapiro-Wilk test of paired differences, P = 0.6142 | Not applicable (paired design) | paired two-tailed t-test | P = 0.0412 (*) | t(9) = -2.3810 |
| Figure 6N<br>Fmr1-KO sociability preference index | n = 10 paired observations | Shapiro-Wilk test of paired differences, P = 0.8876 | Not applicable (paired design) | paired two-tailed t-test | P = 0.0131 (*) | t(9) = -3.0842 |
| Figure 6O<br>WT+Fmr1-I234T sociability preference index | n = 10 paired observations | Shapiro-Wilk test of paired differences, P = 0.1713 | Not applicable (paired design) | paired two-tailed t-test | P = 0.0136 (*) | t(9) = -3.0566 |
| **Supplementary Figure 1** | | | | | | |
| Supplementary Figure 1A<br>Age | FXS = 52;<br>TD = 64 | Shapiro-Wilk test, P(FXS) = 0.0004; P(TD) = 0.0003 | Bartlett's test: P = 0.6235 | Mann–Whitney U test | P = 0.5692 (ns) | U = 1561.0000 |
| Supplementary Figure 1B<br>Sex distribution | FXS: Female = 4, Male = 48<br>TD: Female = 20, Male = 44 | Not applicable | Not applicable | Pearson chi-squared test | P = 0.0018 (**) | $\chi^2(1) = 9.7030$ |
| **Supplementary Figure 3** | | | | | | |
| Supplementary Figure 3A<br>Nose FA | WT = 7;<br>Fmr1-KO = 7 | Shapiro-Wilk test, P(WT) = 0.4182; P(Fmr1-KO) = 0.8852 | Bartlett's test: P = 0.0446 | unpaired two-tailed t-test | P = 0.1777 (ns) | t(12) = 1.4318 |
| Supplementary Figure 3A<br>Barrel FA | WT = 7;<br>Fmr1-KO = 7 | Shapiro-Wilk test, P(WT) = 0.8708; P(Fmr1-KO) = 0.9948 | Bartlett's test: P = 0.9765 | unpaired two-tailed t-test | P = 0.1392 (ns) | t(12) = 1.5837 |
| Supplementary Figure 3A<br>Lower limb FA | WT = 7;<br>Fmr1-KO = 7 | Shapiro-Wilk test, P(WT) = 0.9378; P(Fmr1-KO) = 0.8195 | Bartlett's test: P = 0.6527 | unpaired two-tailed t-test | P = 0.2997 (ns) | t(12) = 1.0839 |

| Figure / Panel and endpoint | Sample size (*n*) | Normality test | Variance test | Analysis | *P* Value | Degrees of freedom and test statistic |
|---|---|---|---|---|---|---|
| Supplementary Figure 3A Trunk FA | WT = 7; Fmr1-KO = 7 | Shapiro-Wilk test, P(WT) = 0.6330; P(Fmr1-KO) = 0.2698 | Bartlett's test: P = 0.5011 | unpaired two-tailed t-test | P = 0.7271 (ns) | t(12) = 0.3573 |
| Supplementary Figure 3B Nose FA | WT = 7; Fmr1-KO = 7 | Shapiro-Wilk test, P(WT) = 0.4194; P(Fmr1-KO) = 0.6620 | Bartlett's test: P = 0.7791 | unpaired two-tailed t-test | P = 0.2453 (ns) | t(12) = 1.2216 |
| Supplementary Figure 3B Barrel FA | WT = 7; Fmr1-KO = 7 | Shapiro-Wilk test, P(WT) = 0.8889; P(Fmr1-KO) = 0.7593 | Bartlett's test: P = 0.5614 | unpaired two-tailed t-test | P = 0.2453 (ns) | t(12) = 1.2215 |
| Supplementary Figure 3B Lower limb FA | WT = 7; Fmr1-KO = 7 | Shapiro-Wilk test, P(WT) = 0.5176; P(Fmr1-KO) = 0.5234 | Bartlett's test: P = 0.4909 | unpaired two-tailed t-test | P = 0.6402 (ns) | t(12) = 0.4795 |
| Supplementary Figure 3B Trunk FA | WT = 7; Fmr1-KO = 7 | Shapiro-Wilk test, P(WT) = 0.5238; P(Fmr1-KO) = 0.4121 | Bartlett's test: P = 0.4544 | unpaired two-tailed t-test | P = 0.4892 (ns) | t(12) = -0.7135 |
| **Supplementary Figure 4** | | | | | | |
| Supplementary Figure 4C Nonsocial-condition AUC | WT = 7; Fmr1-KO = 6 | Shapiro-Wilk test, P(WT) = 0.2312; P(Fmr1-KO) = 0.2474 | Bartlett's test: P = 0.0805 | unpaired two-tailed t-test | P = 0.6862 (ns) | t(11) = 0.4149 |
| Supplementary Figure 4C Nonsocial-condition peak delta-F/F | WT = 7; Fmr1-KO = 6 | Shapiro-Wilk test, P(WT) = 0.6175; P(Fmr1-KO) = 0.0018 | Bartlett's test: P = 0.0188 | Mann–Whitney U test | P = 0.1375 (ns) | U = 10.0000 |
| **Supplementary Figure 5** | | | | | | |
| Supplementary Figure 5D S1-DMS calcium AUC | WT = 11; Fmr1-KO = 9 | Shapiro-Wilk test, P(WT) = 0.2966; P(Fmr1-KO) = 0.4439 | Bartlett's test: P = 0.0002 | Welch's unpaired two-tailed t-test | P = 0.0405 (*) | t(8.8502) = -2.3973 |
| Supplementary Figure 5D S1-DMS calcium peak delta-F/F | WT = 11; Fmr1-KO = 9 | Shapiro-Wilk test, P(WT) = 0.4624; P(Fmr1-KO) = 0.1906 | Bartlett's test: P = 0.1816 | unpaired two-tailed t-test | P = 0.0344 (*) | t(18) = -2.2895 |
| **Supplementary Figure 6** | | | | | | |
| Supplementary Figure 6B Novel location recognition index | Fmr1-KO + Fmr1-WT = 7; Fmr1-KO + Fmr1-I234T = 7 | Shapiro-Wilk test, P(Fmr1-KO + Fmr1-WT) = 0.3818; P(Fmr1-KO + Fmr1-I234T) = 0.5304 | Bartlett's test: P = 0.5205 | unpaired two-tailed t-test | P = 0.0373 (*) | t(12) = 2.3417 |

| Figure / Panel and endpoint | Sample size (*n*) | Normality test | Variance test | Analysis | *P* Value | Degrees of freedom and test statistic |
|---|---|---|---|---|---|---|
| Supplementary Figure 6D Sociability preference index | Fmr1-KO + Fmr1-WT = 8; Fmr1-KO + Fmr1-I234T = 7 | Shapiro-Wilk test, P(Fmr1-KO + Fmr1-WT) = 0.2354; P(Fmr1-KO + Fmr1-I234T) = 0.0836 | Bartlett's test: P = 0.2115 | unpaired two-tailed t-test | P = 0.0007 (***) | t(13) = 4.3790 |
| Supplementary Figure 6F WT + Fmr1-I234T novel location recognition index | n = 12 paired observations | Shapiro-Wilk test of paired differences, P = 0.8347 | Not applicable (paired design) | paired two-tailed t-test | P = 0.0127 (*) | t(11) = -2.9700 |

**Supplementary Table 3. Summary of whole-brain white matter microstructural alterations in FXS compared to TD**

| DTI Metric | Alteration | Cluster Size (Voxels) | Peak MNI Coordinate (X, Y, Z) | Major Anatomical Distributions (JHU Atlas) |
|---|---|---|---|---|
| **FA** | Decrease | 75,120 | 5, -44, -46 | Superficial WM/U-fibers (~66.0%)<br>Corona Radiata (~6.1%)<br>Corpus Callosum (~5.7%)<br>Internal Capsule (~5.3%) |
| | Increase | None | - | - |
| **RD** | Increase | 88,560 | -2, -36, -47 | Superficial WM/U-fibers (~69.4%)<br>Corona Radiata (~5.6%)<br>Corpus Callosum (~5.4%)<br>Internal Capsule (~4.8%) |
| | Decrease | None | - | - |
| **MD** | Increase | 82,444 | -41, -12, -23 | Superficial WM/U-fibers (~69.7%)<br>Corpus Callosum (~7.0%)<br>Corona Radiata (~5.5%)<br>Internal Capsule (~4.5%) |
| | Decrease | None | - | - |
| **AD** | Increase | 27,765 | -18, -46, 8 | Superficial WM/U-fibers (~62.1%)<br>Corpus Callosum (~13.1%)<br>Corona Radiata (~7.5%)<br>Superior Longitudinal Fasciculus (~4.2%) |
| | | 2609 | 2, -55, -25 | Unclassified / Cerebellar WM (~87.4%) |

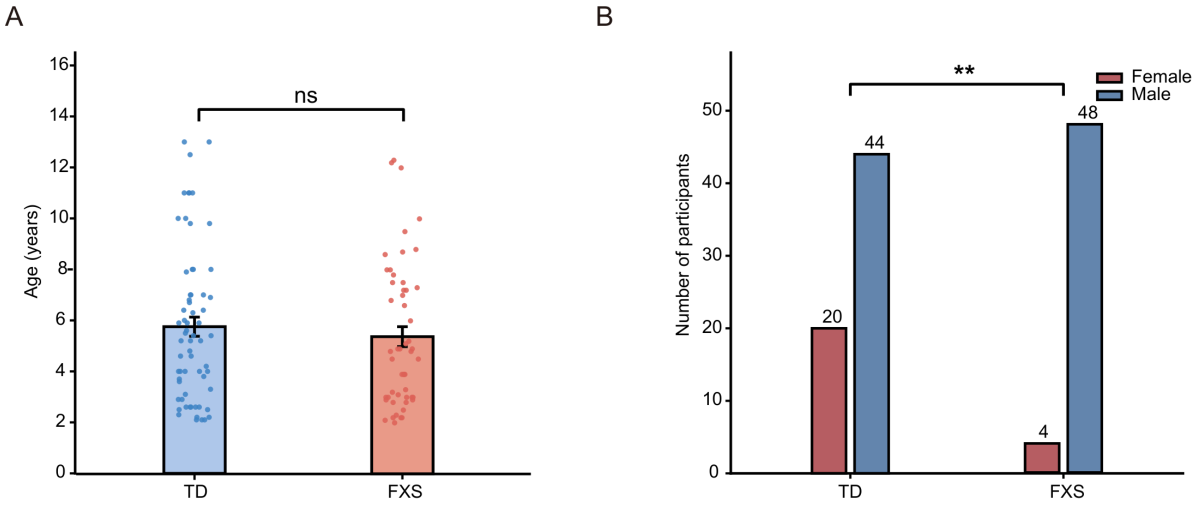


**Supplementary Figure 1. Age and sex distributions in the human cohorts.** (A) Age distributions in participants with fragile X syndrome (FXS; n = 52) and typically developing (TD) participants (n = 64). A two-tailed Mann–Whitney U test was used. (B) Sex distributions. The FXS group comprised 48 male and four female participants, and the TD group comprised 44 male and 20 female participants. Pearson's $\chi^2$ test was used. **P < 0.01; ns, not significant. Age data are presented as mean ± standard error of the mean (SEM); sex data are participant counts.

RD: FXS > TD

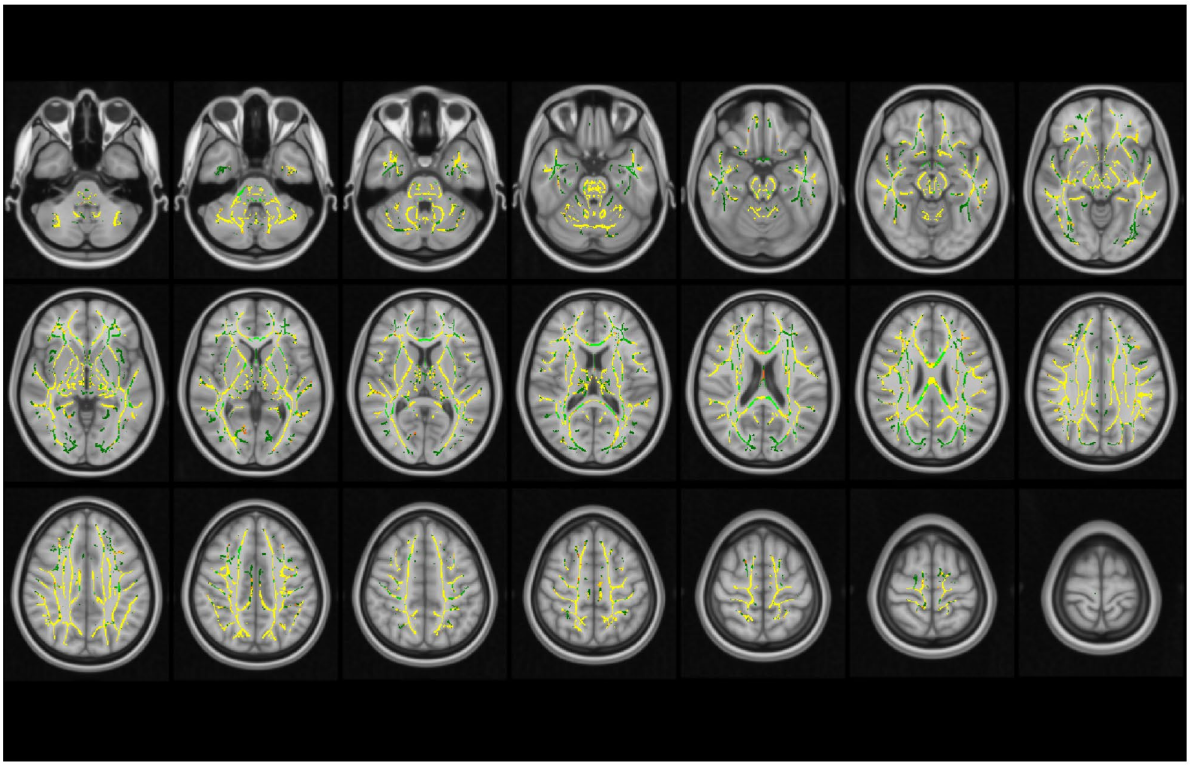

**Supplementary Figure 2. Increased radial diffusivity in participants with fragile X syndrome.** Axial slices show regions with higher radial diffusivity (RD) in participants with fragile X syndrome (FXS; n = 52) than in typically developing (TD) participants (n = 64). Significant voxels are displayed in warm colours over the white-matter skeleton (green) and the MNI152 T1-weighted anatomical template. Tract-based spatial statistics (TBSS) included age and sex as covariates and used threshold-free cluster enhancement (TFCE) with family-wise error (FWE) correction at $P < 0.05$. MNI, Montreal Neurological Institute.

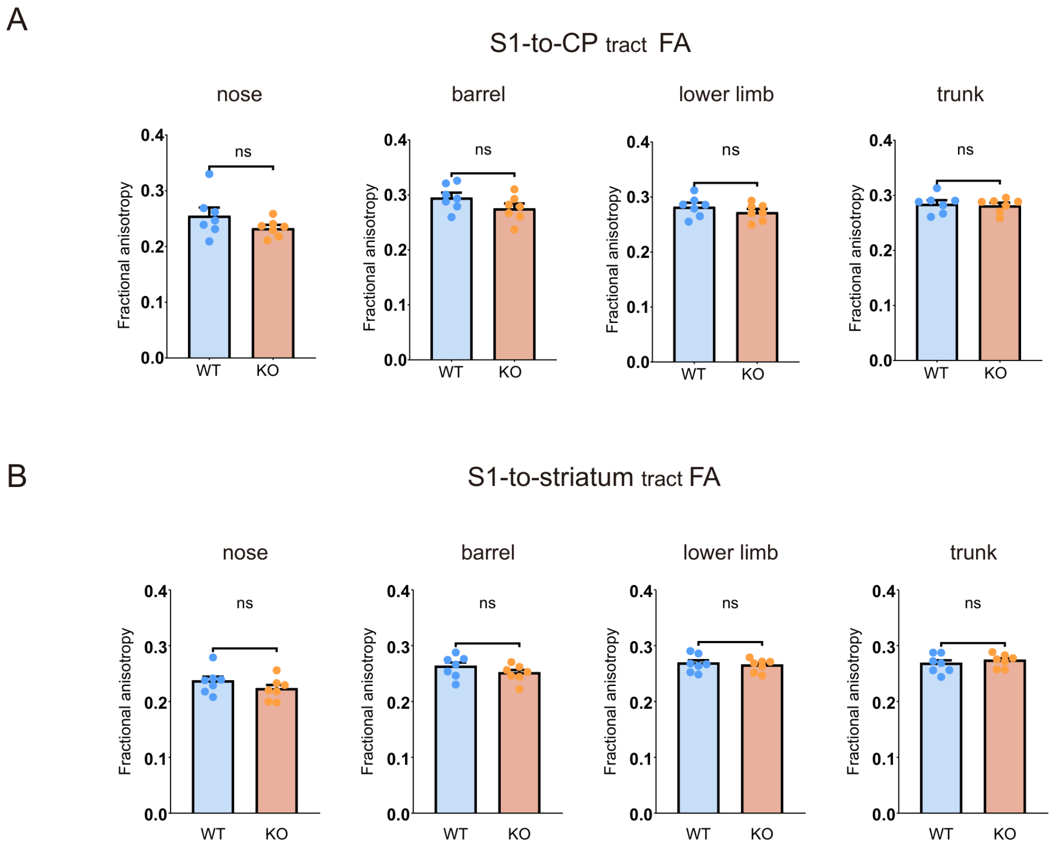


**Supplementary Figure 3. Fractional anisotropy in additional S1–caudoputamen and S1–striatum pathways.** (A) Fractional anisotropy (FA) in pathways connecting the nose, barrel, lower-limb, and trunk subregions of primary somatosensory cortex (S1) to the caudoputamen (CP), from left to right. (B) FA in the corresponding S1-to-striatum pathways, in the same subregion order. Wild-type (WT) and *Fmr1*-knockout (KO) mice, n = 7 per genotype. Unpaired two-tailed t-tests were used for all comparisons in A and B. Individual points represent mice. Data are presented as mean ± SEM. ns, not significant.

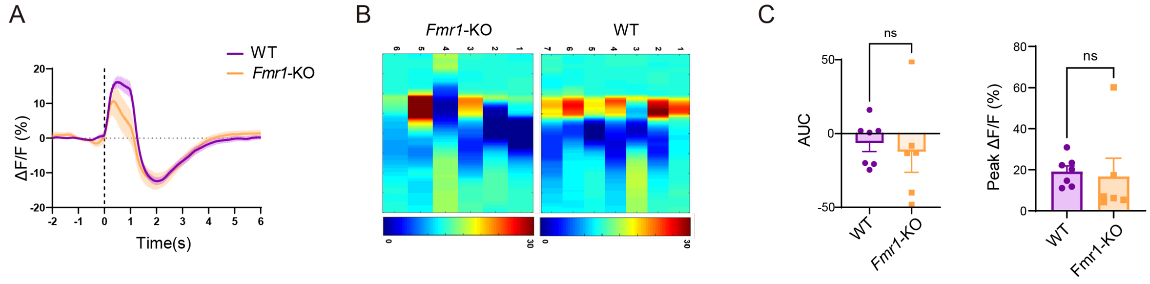


**Supplementary Figure 4. S1 calcium responses to local optogenetic stimulation in a nonsocial condition.** (A–C) Peri-event mean calcium traces (A), animal-level heat maps (B), and response area under the curve (AUC) and peak ΔF/F (C) during a nonsocial condition with local primary somatosensory cortex (S1) stimulation. S1 stimulation and recording used CaMKIIα-ChrimsonR and CaMKIIα-GCaMP6m. Wild-type (WT), n = 7 mice; *Fmr1*-knockout (KO), n = 6 mice. An unpaired two-tailed t-test was used for AUC and a two-tailed Mann–Whitney U test for peak ΔF/F. ns, not significant. Data are presented as mean ± SEM.

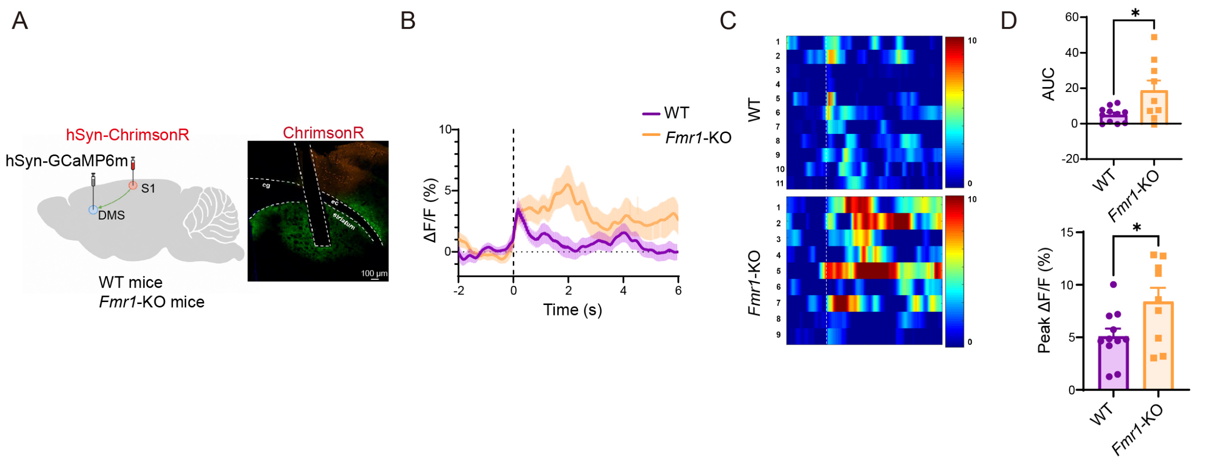


**Supplementary Figure 5. DMS calcium responses to S1 optogenetic stimulation in *Fmr1*-deficient mice.** (A) Configuration for ChrimsonR-mediated primary somatosensory cortex (S1) stimulation and dorsomedial striatum (DMS) calcium recording using hSyn-GCaMP6m, with representative histology (scale bar, 100 µm). (B–D) Peri-event mean calcium traces (B), animal-level heat maps (C), and response area under the curve (AUC) and peak ΔF/F (D). Wild-type (WT), n = 11 mice; *Fmr1*-knockout (KO), n = 9 mice. A two-tailed Welch t-test was used for AUC (P = 0.0405), and an unpaired two-tailed t-test for peak ΔF/F. *P < 0.05. Data are presented as mean ± SEM.

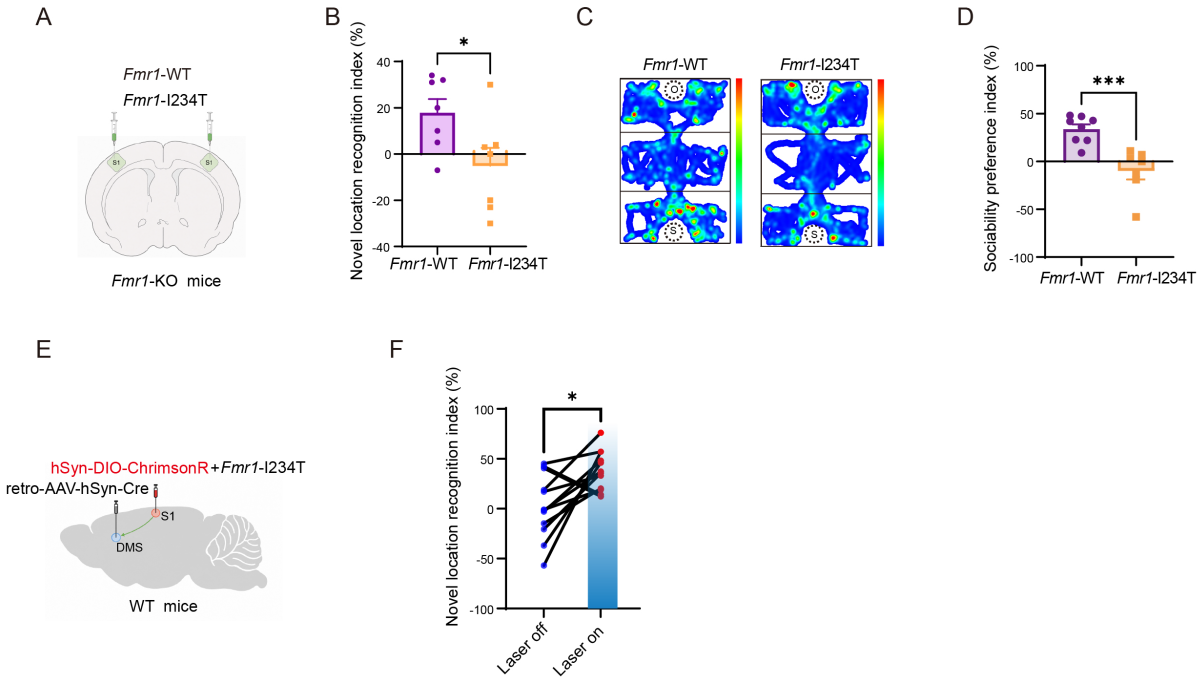


**Supplementary Figure 6. *Fmr1*-I234T expression and S1–DMS projection activation modify behavioural phenotypes.** (A) Local expression of *Fmr1*-WT or *Fmr1*-I234T in primary somatosensory cortex (S1) of *Fmr1*-knockout (KO) mice. (B) Novel-location-recognition indices (n = 7 mice per group). (C and D) Representative sociability heat maps (C) and preference indices (D; *Fmr1*-KO + *Fmr1*-WT, n = 8 mice; *Fmr1*-KO + *Fmr1*-I234T, n = 7 mice). (E) Activation of dorsomedial striatum (DMS)-projecting S1 neurons during novel-location-recognition testing in wild-type (WT) mice expressing *Fmr1*-I234T in S1, using retrograde Cre delivery to DMS and Cre-dependent ChrimsonR expression in S1. (F) Paired novel-location-recognition indices under laser-off and laser-on conditions (n = 12 mice). Unpaired two-tailed t-tests were used for B and D, and a two-tailed paired t-test for F. *P < 0.05, ***P < 0.001. Data are presented as mean ± SEM.

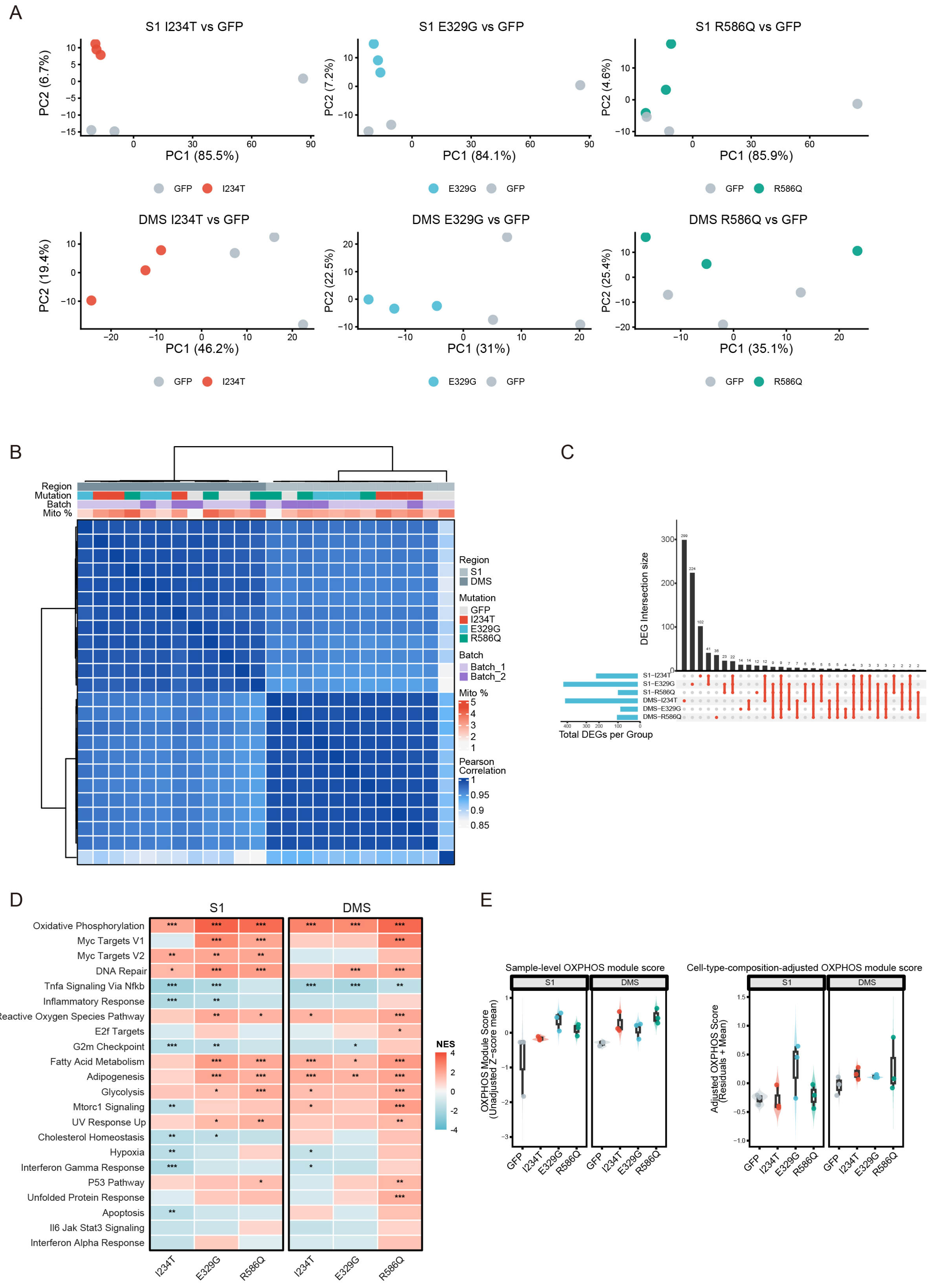
A
S1 I234T vs GFP
S1 E329G vs GFP
S1 R586Q vs GFP
PC1 (85.5%)
PC2 (6.7%)
PC1 (84.1%)
PC2 (7.2%)
PC1 (85.9%)
PC2 (4.6%)
GFP I234T
E329G GFP
GFP R586Q
DMS I234T vs GFP
DMS E329G vs GFP
DMS R586Q vs GFP
PC1 (46.2%)
PC2 (19.4%)
PC1 (31%)
PC2 (22.5%)
PC1 (35.1%)
PC2 (25.4%)
B
Region
Mutation
Batch
Mito %
Region
S1
DMS
Mutation
GFP
I234T
E329G
R586Q
Batch
Batch_1
Batch_2
Mito %
Pearson
Correlation
C
DEG Intersection size
S1−I234T
S1−E329G
S1−R586Q
DMS−I234T
DMS−E329G
DMS−R586Q
Total DEGs per Group
D
S1
DMS
Oxidative Phosphorylation
Myc Targets V1
Myc Targets V2
DNA Repair
Tnfa Signaling Via Nfkb
Inflammatory Response
Reactive Oxygen Species Pathway
E2f Targets
G2m Checkpoint
Fatty Acid Metabolism
Adipogenesis
Glycolysis
Mtorc1 Signaling
UV Response Up
Cholesterol Homeostasis
Hypoxia
Interferon Gamma Response
P53 Pathway
Unfolded Protein Response
Apoptosis
Il6 Jak Stat3 Signaling
Interferon Alpha Response
I234T
E329G
R586Q
NES
E
Sample-level OXPHOS module score
Cell-type-composition-adjusted OXPHOS module score
S1
DMS
OXPHOS Module Score
(Unadjusted Z-score mean)
Adjusted OXPHOS Score
(Residuals + Mean)
GFP
I234T
E329G
R586Q

**Supplementary Figure 7. Transcriptomic structure and OXPHOS pathway scores from regional RNA-sequencing analysis.** (A) Principal-component analysis (PCA) of each *Fmr1*-variant comparison with GFP in primary somatosensory cortex (S1) and dorsomedial striatum (DMS). (B) Pearson sample-correlation heat map with annotations for region, construct, batch and percentage of mitochondrial reads. (C) UpSet plot of intersections among differentially expressed gene (DEG) sets. (D) Normalised enrichment scores (NESs) for Hallmark gene sets across the six region-by-variant comparisons. (E) Sample-level oxidative phosphorylation (OXPHOS) module scores before and after adjustment for cell-type-marker composition. Each region-by-construct group comprised three independent bulk-tissue RNA-sequencing libraries, with 24 libraries in total. DEGs were defined using DESeq2 with adjusted $P < 0.05$ and $|\log 2$ fold change$| \geq \log 2(1.5)$. PC1 and PC2 denote the first and second principal components; GFP, green fluorescent protein; VST, variance-stabilising transformation; Mito %, percentage of mitochondrial reads. Gene-set enrichment analysis (GSEA) used false discovery rate (FDR) correction. In D, *FDR $q < 0.05$, **FDR $q < 0.01$, ***FDR $q < 0.001$.
